# Joint modeling of choices and reaction times based on Bayesian contextual behavioral control

**DOI:** 10.1101/2021.10.29.466505

**Authors:** Sarah Schwöbel, Dimitrije Markovic, Michael N. Smolka, Stefan Kiebel

## Abstract

In cognitive neuroscience and psychology, reaction times are an important behavioral measure. However, in instrumental learning and goal-directed decision making experiments, findings often rely only on choice probabilities from a value-based model, instead of reaction times. Recent advancements have shown that it is possible to connect value-based decision models with reaction time models. However, typically these models do not provide an integrated account of both value-based choices and reaction times, but simply link two types of models. Here, we propose a novel integrative joint model of both choices and reaction times by combining a mechanistic account of Bayesian sequential decision making with a sampling procedure. This allows us to describe how internal uncertainty in the planning process shapes reaction time distributions. Specifically, we use a recent context-specific Bayesian forward planning model which we extend by a Markov chain Monte Carlo (MCMC) sampler to obtain both choices and reaction times. As we will show this makes the sampler an integral part of the decision making process and enables us to reproduce, using simulations, well-known experimental findings in value based-decision making as well as classical inhibition and switching tasks. Specifically, we use the proposed model to explain both choice behavior and reaction times in instrumental learning and automatized behavior, in the Eriksen flanker task and in task switching. These findings show that the proposed joint behavioral model may describe common underlying processes in these different decision making paradigms.

**Author summary:** Many influential results in psychology and cognitive neuroscience rest on reaction time effects in behavioral experiments, for example in studies about human decision making. For decisions that rest on planning, findings often rely on analyses using specific computational models. Until recently, these models did not allow for analysis of reaction times. In this article we introduce a new model of how to explain both choices and reaction times in decision making experiments that involve planning. Importantly, the model explains how the brain can make good decisions quickly, even in the face of many potential choices and in complex environments.

## Introduction

Many key findings in psychology and cognitive neuroscience are based on the measurement and analysis of both response accuracy and reaction times in behavioral experiments. For example, changes in both mean reaction times and response accuracy during and after conflicting decisions are typically interpreted to demonstrate underlying decision making processes. Such effects of classical experimental paradigms are remarkably stable and have also been used to show how decision making and cognitive control is impaired in several mental disorders [1, 2, 3, 4, 5, 6, 7].

However, these classical experiments typically do not consider the influence of state uncertainty and reward structure on the decision making process. These influences are often investigated using sequential decision-making tasks, as seen in the instrumental learning and value-based decision making literature, e.g. [8, 9, 10]. Still, in these research areas, decision making under uncertainty is mostly understood by applying value-based decision models using associated choice probabilities and model parameters. With value-based decision making models, reaction time effects are rarely explicitly modeled and investigated. A likely reason is that the considered computational behavioral models are based on research in reinforcement learning and Bayesian decision making, which is not primarily aimed at describing reaction times associated with decisions.

To fill this gap, there have been recent, successful combined applications of value-based decision and reaction time models, specifically by coupling reinforcement learning with a reaction time model [11, 12, 13, 14, 15, 16], which are typically evidence accumulator models such as drift diffusion models (DDM) [17, 18] or so-called race diffusion models [11]. The principled idea is to connect choice values and probabilities to reaction times by linking parameters in both models. For example, the trial-wise expected reward (Q-values) in reinforcement learning models has been used to vary the drift rate of a DDM [13]. This approach has been extended to multi-choice tasks using race diffusion models, where instead of having one accumulator as in the DDM, each available choice option is associated with a different accumulator [14, 16].

Although this type of coupled model is useful to analyze reaction times and choices jointly, the coupling between the two models is unidirectional. Specifically, the reinforcement learning model computes expected rewards in isolation and feeds its output forward to the DDM, which adds noise to the decision by sampling, thereby returning RT and choice distributions [12]. An alternative role for sampling processes in decision making has been proposed recently under the Bayesian brain hypothesis, specifically active inference. In this view, sampling, here Monte Carlo sampling, takes an integral role in the computation of Bayesian approximate inference in order to model cognitive processes as for example evidence accumulation [19, 20, 21]. In comparison to standard reinforcement learning models, such as Q-learning, a Bayesian approach makes it explicit how the brain forms beliefs based on internal representations and deals with uncertainty in observations and planning. With such an approach, one can view measured reaction time distributions as a direct expression of planning under uncertainty. This may enable the analysis of planning parameters which would not be accessible with a forward-coupled model, e.g. RL and DDM. For example, it has been proposed that reaction times are related to information processing and encoding costs [22], and may relate to cognitive effort in active inference [22, 23].

Within this Bayesian active inference perspective, we propose an integrated choice and sampling reaction time model, the Bayesian contextual control (BCC) model, where reaction times are determined by the encoding of information and planning process itself. To model classic psychological experiments, we will use a recently introduced Bayesian model of context-dependent behavior as a basis [24]. The model quantitatively describes forward planning and goal-directed decision making, as well as repetition-based choice biases, in a context-dependent way. This will allow us to model switching between task sets (as in a task switching task), as well as automatic behavioral biases (as in a flanker task). Into this BCC model, we integrate an independent Markov chain Monte Carlo (MCMC) sampler to mechanistically describe how choices and reaction times emerge from the internal planning process. Reaction times in this model depend on the number of possible options, the uncertainty in the planning process, as well as conflicts between contexts or between goal-directed actions and biases. As we will show below, this new approach allows us to account for four key components of behavioral effects: (i) the typical log-normal shapes of reaction time distributions, (ii) uncertainty at multiple levels affecting reaction times, (iii) biases for cued response repetitions, and (iv) context-specific effects.

To illustrate the properties of the new model, we will use two toy examples. We show how the sampling works internally, that the model can replicate the classical log-normal reaction time distributions, and that the strength of prior beliefs decrease reaction times and error rates.

Furthermore, to demonstrate the versatility of the model, we will show simulated behavior of two widely used behavioral experiments: the Eriksen flanker task, and a task switching task, which are often used to study cognitive processes and impaired cognitive control [1, 2]. In the flanker task, conflicting information in incongruent trials leads to longer reaction times due to a goal-bias conflict which induces uncertainty on the action selection. In addition, due to the sampling process, responses with shorter reaction times exhibit higher error rates, as is typically found in the flanker literature. In the task switching task, training reduces reaction times and error rates, due to decreasing uncertainty about the task structure. Switching between task sets, which are interpreted as contexts in the model, increases reaction times and error rates due to a conflict between contexts, which introduces uncertainty into the planning process. In both simulated tasks, we can replicate well-known effects commonly reported in the literature. We close by discussing the implications of the underlying mechanism and its relation to alternative models.

## Methods

In this section, we briefly describe the computational model of behavior used for simulations, and the novel reaction time component which was used to generate agent’s reaction times.

### Bayesian contextual control model

Here, we will outline the Bayesian contextual control (BCC) model and key aspects that influence reaction times and the underlying mechanisms. The full mathematical details of the BCC model are provided in the appendix Supplementary 1: Detailed mathematical derivations and a Python implementation is publicly available on github^1^.

The approach is based on the ideas of planning as inference [25, 26], active inference [27, 28, 29, 30], and our previous work [24], where the dynamics of the environment, as well as the outcome contingencies are represented in a Bayesian generative model. This model is used to evaluate a posterior over actions, or action sequences, from which actions are chosen. In line with the active inference literature, we will hereafter call a deterministic action sequence a policy *π* [31, 30].

As in our previous work [24], we aim at modeling an agent that can switch adaptively between different environmental contexts, e.g. task conditions. This model structure has the advantage that an agent can switch between context-specific policies guided by a context-specific task representation and a learned prior over policies, thereby emulating the participants’ switching between task conditions. The key idea of this paper is to use a sampling mechanism to evaluate the posterior over these context-specific policies to obtain reaction times to create an integrated models of choices and reaction times.

In the model, see Fig 1 for a graphical representation, the dynamics of the environment are clustered into behavioral episodes, where an episode in our model corresponds to the length of the task at hand. In most experiments, this would translate into one trial, and in sequential decision tasks, to the length of the sequence. Within an episode, the state transitions and outcome contingencies are determined by the current context, e.g. the current task condition. The context-specific representation of the task at hand in the form of a Markov decision process (MDP) is updated and learned after each episode that is experienced in a context. The context itself is treated as a hidden variable, that is inferred based on cues or experienced contingencies.

**Fig 1.**
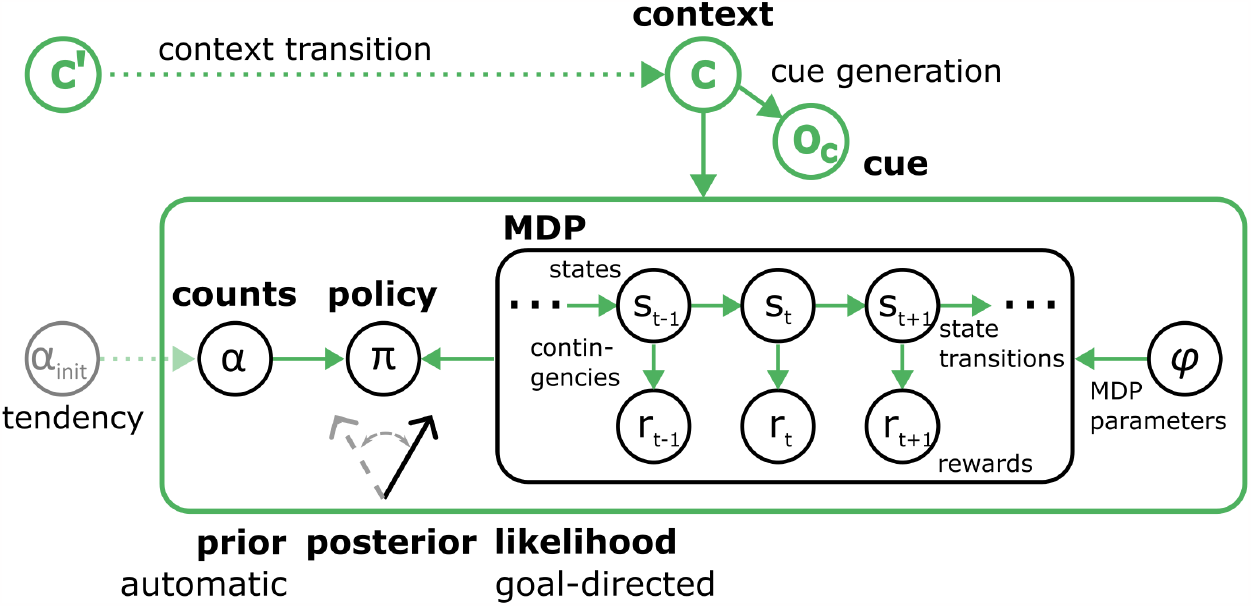
Graphical sketch of the generative model. Circles depict variables in the generative model, while arrows show conditional dependencies, and the green color indicates context-dependent parts or parameters of the model. The inner black box shows the goal-directed component of the model in the form of an MDP, where states (e.g. *s*_*t*_) evolve over time according to state transition rules (right pointing arrows). Depending on the states, rewards (*r*_*t*_) may be generated in each time step according to outcome contingencies (down arrows) and the current state, where both may be context dependent. Values of state transition rules and contingencies are treated as hidden variables *ϕ* (black circle on the right) in the model as well, and are updated based on experience, which enables learning. This MDP is of fixed length and constitutes an episode. State transitions and rewards also depend on the behavioral policy *π* (circle left of the box). This MDP is used to calculate the likelihood *p*(*R*|*π, c*) of receiving rewards under a policy. The left most circle in the green box (*α*) represents the hyper parameters of the prior over policies *p*(*π*|*c, α*), which encodes automatic a priori biases for the policies. The initial parameters *α*_*init*_ (grayed out box left of the green box) are free parameters that can represent initial behavioral biases when a context is encountered for the first time. For a detailed description of the free parameters and their relation to behavior and experiments, see Section Free parameters and simulation setups. Prior and likelihood are used to calculate the posterior over policies p(*π*|*R, c*), see Eq. 1. The outer green box indicates all components within the box are context-dependent. The context *c*, green circle on the top, is a hidden variable that determines the parameter values that are used for the prior and likelihood. The context needs to be inferred using available information.

This can be sketched in a Bayesian equation as

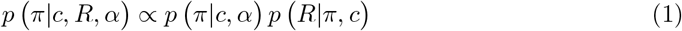

which describes the posterior probability *p*(*π*|*c, R, α*) of whether an agent should choose policy *π*, given desired rewards *R* in the current context *c*, see Fig 1. This posterior is a categorical distribution and according to Bayes’ rule it is proportional to the likelihood of obtaining rewards under a policy in the current context *p*(*R*|*π, c*) times a context-specific prior over policies *p*(*π*|*c, α*).

Importantly, the likelihood of rewards *p*(*R*|*π, c*) is calculated based on the current model (i.e., an MDP), which contains outcome contingencies and therewith encodes a goal-directed value of a policy. We use variational inference to calculate beliefs over future states and rewards, which are then averaged out to obtain the expected reward under a policy. The prior over policies (*p*|*π c, α*) on the other hand, is based on counts *α*, which are updated and learned based on Bayesian learning rules. This update mechanism yields higher a priori probabilities for policies which have been previously chosen in this context. We interpret this prior updating as repetition-based automatism learning, because this term implements a priori biases to repeat policies, independent of any reward expectations [24]. Due to the prior being over policies, i.e. full action sequences, this implements context-dependent automatism biases for certain action sequences, where behavior can be a continuum from fully goal-directed (no bias) to fully automatic (extreme prior for one specific sequence). On the intermediate points on this continuum, the prior acts as a heuristic that guides behavior in well-known contexts. The posterior natively balances the automatic and goal-directed behavioral contributions using Bayes’ theorem.

Response conflicts can emerge when the context is not directly observable, and there is uncertainty over the current context that cannot be fully resolved. Hence, the agent may not know with certainty which rules of the environment currently apply. To enable context inference, we introduced context observations *o*_*c*_, i.e. cues, into the model, and the agent maintains cue-generation probabilities for specific contexts *p*(*o*_*c*_|*c*). Beliefs over contexts can also be inferred through the experience in an episode, e.g. the experienced outcome contingencies. Additionally, the beliefs over the context depend on the beliefs over the previous context *c*′, which are propagated through context transition probabilities *p*(*c*|*c*′). Using Bayesian inversion, the agent can infer a posterior over contexts *p*(*c*|*o*_*c*_).

The resulting posterior over policies

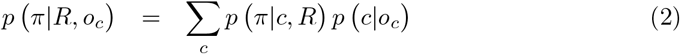

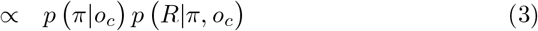

is a mix of the context-specific posteriors over policies, weighted by the posterior probabilities of each context. The posterior over policies *p*(*π*|*R*) gives the probability that an agent should choose a specific policy *π*, given it wants to receive rewards *R*. This posterior is proportional to the prior over policies *p*(*π*) times the likelihood of receiving rewards *p*(*R*|*π*). The behavioral policy that is actually executed is selected from this posterior and the concrete procedure which implements the planning process is described below. In a conflict situation, the two conflicting policies would be similarly weighted in the posterior, which leads to an increased error rate, as they would be similarly likely to be selected.

Given this basis, we will now describe the sampling process, which will give rise to reaction times.

### Planning and reaction times

The key idea is that our model agent not only learns and recalls context-specific priors over policies as a behavioral heuristic for a given context, but also uses these to guide the planning process itself. This allows for fast action selection in familiar, well-learned situations, or when there is a tight deadline for selecting the action. The goal-directed likelihood of receiving rewards *p*(*R*|*π*), is costly to evaluate fully, as it requires forward propagation of beliefs over (latent) states, and therefore an exhausting planning process is computationally slow. Instead, we propose that the prior is used to iteratively sample policies for which the likelihood is then evaluated, to implement a priority-based planning procedure. During planning, the posterior *p*(*π*|*R*) is iteratively approximated by a sampling-based distribution *q*(*π*). Sampling and planning conclude once the agent is sufficiently certain to have sampled enough policies (under a given time pressure limit) for a satisfactory estimate of the posterior *q*(*π*) *≈ p*(*π*|*R*). This certainty-based sampling duration is, from the view of an agent, a time-saving process for two reasons: (i) well-learned context-specific priors over policies can be very precise, so that the agent would only sample tightly around few or even a single policy; and (ii) under time pressure, promising policies will be sampled first, which allows for fast but accurate planning. This process is shown in Fig 2A. Note that the sampling process makes this planning process noisy, as the order of sampled policies may vary, which adds noise to the reaction times leading to the classical reaction time distributions, which we show below (see Section Value-based decision making in a grid world). Importantly, this means that the noise in this model is a direct effect of the reward prediction and action selection processes, and an integral part of the planning process.

**Fig 2.**
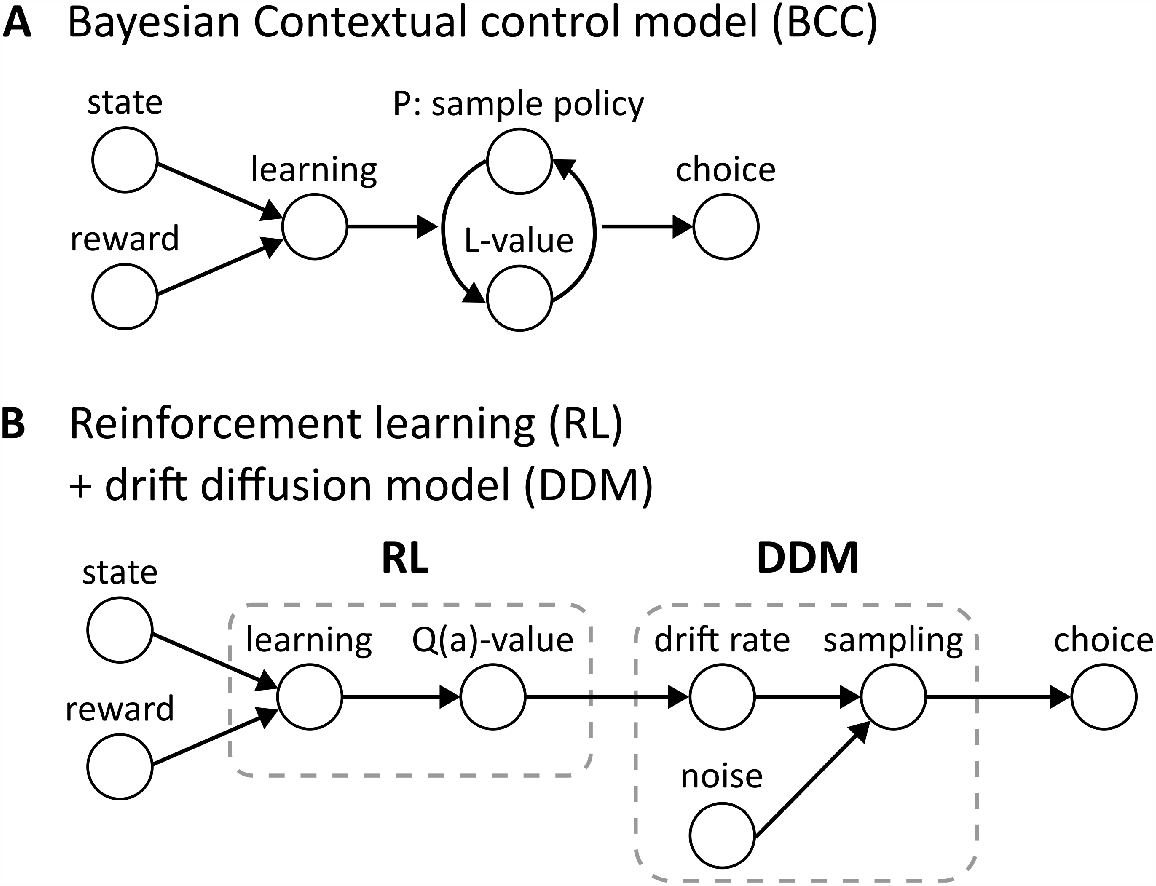
Mechanisms of choice and RT generation in the BCC and the RL+DDM models. (A) The BCC model uses experienced states and rewards as inputs, which update the underlying MDP, as well as parameters and priors through learning. To generate a choice, a policy is sampled from the prior over policies *P* = *p*(*π*) and expected rewards under this policy are evaluated through the likelihood *L* = *p*(*R*|*π*). This process is iterated until a stopping criterion is reached, and the agent is sufficiently certain that the sampling approximates the true posterior *q*(*π*) *≈ p*(*π*|*R*). The action according to the most recently sampled policy is then chosen and executed. Variability in RTs and choices is a direct consequence of the policy sampling for the calculation of the predicted rewards, and therewith an intrinsic property of the planning process in the BCC model. (B) A typical RL+DDM value-based decision making and RT model uses states and rewards as inputs to an instance of an RL model. These are used to update the underlying model through learning, and calculate reward-based action values (Q-values), which are used to select actions. The action values are fed into an instance of a DDM, or more generally, an evidence accumulator model. The action values are used to set DDM parameters, such as the drift rate. In each sampling step, Gaussian white noise is added to introduce variability to the sampled choices and RTs. Finally, when the sampling reaches a threshold, a choice is selected and executed.

In summary, we propose here a mechanism of how reaction times emerge from the planning process itself. The resulting reaction time is influenced by the integral noise of the sampling process, stemming from the uncertainty of actions and outcomes, and the similarity of prior and likelihood. This is a fundamentally different approach to the standard combined RL+DDM approach, as shown in Fig 2B, where internal planning parameters like Q-values are being calculated in isolation in the value-based part of the model, and are then passed on to the DDM. The noise is then added in the DDM as Gaussian noise, which is independent from the planning process in the RL model.

However, even if some aspects are not modeled explicitly, the decades of literature on the DDM clearly show that it provides a good description of reaction time generation. Additionally, the DDM is a general sampling model that can describe many decision making modes. In this paper, our main goal is to present a novel and interesting potential mechanism for reaction time generation based on planning and context uncertainty, rather than to propose a general alternative to the DDM.

Mathematically, this process can be described using Markov chain Monte Carlo (MCMC) methods, specifically a modified independent Metropolis-Hastings algorithm which yields a Bayesian independence sampler. Each sampling step can be described with the following equations:

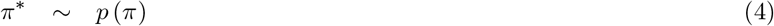

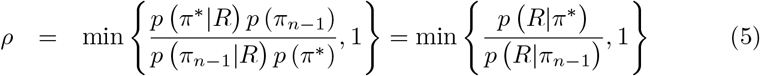

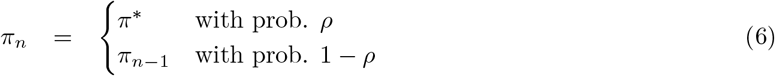

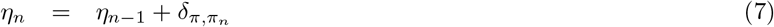

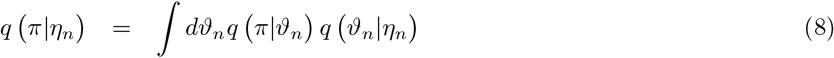

A sample policy *π*^*\**^ is drawn from the prior over policies *p* (*π*) (Eq 4), after which the likelihood *p*(*R*|*π*^*\**^) of the sample *π*^*\**^ is evaluated, and the sample is accepted or rejected into the chain based on the ratio *ρ* of the likelihoods of the current and the previous sample (Eq 5, Eq 6). When the chain has converged, the samples in the chain constitute i.i.d. drawn samples from the posterior over policies. This allows an agent to estimate the posterior over policies through the samples in the chain.

Since the posterior being estimated through sampling is a categorical distribution, we can infer the parameters *ϑ*_*n*_ of the approximated distribution *q(π*|*ϑ*_*n*_) from the entries of the chain, using a Dirichlet prior *q(ϑ*_*n*_|*η*_*n*_). In each sampling step, the pseudo count of the Dirichlet prior *η*_*n*_ is increased for the policy accepted into the chain (Eq 7). This allows for an online updating of the approximated posterior *q(π*|*η*_*n*_) based on the current pseudo counts (Eq 8). In the next section, we describe how the sampling process translates into reaction times.

#### Certainty-based stopping criterion

To model reaction times, we propose that the sampling concludes once a sufficient level of certainty about the distribution is reached. To achieve this we use the Dirichlet distribution *q(ϑ*|*η*), whose entropy encodes how certain one can be of having found the true parameters, i.e. *q(π*|*η) ≈ p*(*π*|*R*), where a lower entropy corresponds to more certainty. Hence we use a threshold value *H*_*thr*_ = *H*_*init*_ + (*H*_*init*_ *−* 1) *\* s* of the Dirichlet entropy *H[p*(*θ*|*η*)] as a stopping criterion for the sampling, where the free parameter s *∈* (0, *∞*) relates the threshold value to the initial entropy *H*_*init*_. The parameter *s* determines how much lower than the initial entropy the current entropy has to become before sampling concludes, and additionally implements a constant value, in case the initial entropy is 0. This way, the parameter *s* will allow us to up- and down-regulate sampling duration and thereby reaction times (see Section Value-based decision making in a grid world for an illustration of its influence on reaction times). We use the initial entropy, since the entropy may become negative for continuous distributions, such as the Dirichlet distribution, to define a relative threshold value. Note that *s* may have different absolute values for different numbers of policies.

Once the entropy has fallen below this threshold, the last sample in the chain determines which policy is executed. The number of samples the chain *N*_*samples*_ required before finishing is taken as an analogue of the reaction time. Additionally, the shapes of the input distributions influence mean reaction times. Intuitively, under this algorithm sampling concludes earlier and reaction times are faster, the more pronounced prior, likelihood, or the corresponding posterior over policies are. Vice versa, for wide distributions representing uncertainty, e.g. when the goal is unknown or uncertain, sampling continues longer and reaction times become slower. Specifically, reaction times will be slower, when prior and likelihood are in conflict because the sampling process must sample longer to resolve this conflict.

#### Generation of reaction times from sampling steps

To describe measured reaction time data, we assume that the true reaction time in milliseconds is linearly related to the number of samples by

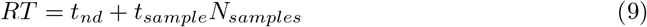

multiplying the number of samples *N*_*samples*_ with a sampling time *t*_*sample*_, and adding a non-decision time, as usually done with DDMs, *t*_*nd*_ which is due to perceptual processes and loading of information [32]. In the simulations of the flanker and task switching tasks below, we set *t*_*sample*_ = 0.2*ms* and *t*_*nd*_ = 100*ms*, to map the number of samples to reaction times below 1000*ms*.

### Free parameters and simulation setups

The resulting model has four free parameters which we will vary in the Results section to recreate experimentally established reaction time effects in behavioral paradigms (see also Supplementary 1: Detailed mathematical derivations). To simulate the different tasks, we adjust the four free parameters, which either reflect the task setup or trait variables: (i) automatization tendency: hyper-parameters of the prior over policies which determine how strong the prior is preset or learned. This would translate to a trait. (ii) The context transition probability: how strongly an agent holds on to the current context. This can be applied to inter-trial effects. (iii) The context cue uncertainty: how well a context cue is perceived. This can describe cue presentation time effects. (iv) Speed-accuracy trade-off: the parameter *s* in the sampling stopping criterion, which decides how long and accurate sampling should be. In this section we will describe the four free parameters and how they relate to experimental setups.

Note that the machinery the agent uses for inference and planning is the same in all the tasks shown in the Results section, as depicted in Fig 1. This means that the process of perceiving a context cue, inferring the context, loading the specific action–outcome contingencies and prior, planning ahead and then sampling an action are common to all setups and all simulations below.

#### Automatization tendency

For learning the prior over policies, the values in the prior are treated as latent random variables (hyper-priors) that follow a Dirichlet distribution. The Dirichlet distribution is parameterized using concentration parameters or pseudo counts *α* of how often a policy was chosen in a specific context, enabling repetition-based learning. While the counting rule is given by the Bayesian updates, the initial values from which counting starts are free parameters which can be chosen at the beginning of a simulation, see also Fig 1. We defined an automatization tendency 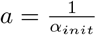, where *a* = 1.0 means the counting starts at 1, giving each new choice a strong influence on the prior over policies. Hence we term an agent with high automatization tendency an “automatism+value learner”. Lower values of *a*, e.g. *a* = 0.001, mean the counting starts at high values, e.g. *α* = 1000, which has the effect that each new choice has little influence on the prior over policies. We call such an agent a “value-only learner”, as in this setting the prior learning is almost negligible as the pseudo counts are dominated by initial values. We show the effects of automatism+value and value-only learning on reaction times and accuracy in a sequential decision task (see Section Value-based decision making in a grid world). Additionally, the pseudo counts can be subjected to a forgetting factor, which takes a similar role to a learning rate (see Supplementary 1: Detailed mathematical derivations) and has them slowly decrease over the course of an experiment, so that later choices still have a measurable effect on the prior over policies. In a stable environment (in most of the simulations shown here), the forgetting factor can be zero, but it is required in dynamically changing environments.

Lastly, not all initial pseudo counts need to be set to the same values. In order to model a priori context–response associations, the initial values can be set so that the prior over policies initially has a bias for specific actions or policies in a specific context. We use such a priori context–response associations to model interference effects in the Flanker task (see Section Flanker task).

#### Context transition probability

The free parameter of the context transition probability encodes an agent’s assumption of how likely a context change is expected to occur after the end of a behavioral episode, which makes it easier or harder for an agent to switch to a new context, see also Fig 1. For example in a task switching experiment, two task sets would correspond to two contexts, and the context transition probability encodes how often an agent thinks the current task set will change. For example, with a low value, an agent expects to stay within the same context, and even a cue to the contrary may not be enough for the agent to infer that the context changed. Traces of the previous context may then still be present even after a context change. If set high, an agent expects a context change to happen after every episode which makes inference of an actual context change more likely, but may also lead to an agent falsely inferring a context change. We will use this effect to model inter-trial effects in a task switching experiment (see Section Task switching).

#### Context cue uncertainty

The context cue uncertainty encodes how certain an agent is about having perceived a specific context cue, see also Fig 1. For example in a task switching experiment, the current task set is cued, and the uncertainty determines how well an agent perceived the cue. A high uncertainty means an agent may not always rely on the cue and may more rely on the previous context to make decisions, while a low uncertainty means an agent perceived the cue well and can reliably infer being in the cued context. We use this context cue uncertainty to model known cue presentation time effects on reaction times in a task switching experiment (see Section Task switching).

#### Speed-accuracy trade-off

The factor *s* in the stopping criterion of the reaction time algorithm implements a speed-accuracy trade-off, and determines when the sampled estimate of the posterior is “good enough”. Larger values lead to longer sampling, making the estimation of the posterior over policies, and the resulting choice of behavior, more accurate while taking longer. Low values of *s* mean that sampling is stopped rather quickly. Importantly, another effect of this speed-accuracy trade-off in the proposed sampling algorithm is that for lower *s* the choice is more likely to adhere to the prior, and less to the not yet accurately estimated likelihood. This means that fast choices will tend to rely on policies with a high prior, which may be interpreted as automatic behavior. We will show these effects in more detail in Section Sampling trajectories in the reaction time algorithm.

## Results

We will first show properties of the BCC model using simulations. To keep these simulations deliberately simple we let agents learn paths to goals in a so-called grid world environment, as used before in theoretical neuroscience, e.g. [33, 34, 35]. Using this environment, we show choice behavior and reaction time effects during learning in the grid world sequential decision task. We go on to show exemplary sampling trajectories and RT distributions to illustrate the properties of the sampling mechanism, including the speed-accuracy trade-off during decision making.

Secondly, we show that the BCC model can also qualitatively explain reaction time changes in standard experimental cognitive control tasks. To do this we adapt the three parameters of the model that reflect differences in experimental setups: The automatization tendency, the context transition probability, and the context cue uncertainty (see Section Free parameters and simulation setups). We use two different tasks: (i) the Eriksen flanker task which is typically interpreted to measure the inhibition aspect of cognitive control, and (ii) Task Switching, which is usually interpreted to measure the cost of switching.

### Value-based decision making in a grid world

In this section, we present properties of the BCC model, and show that we can replicate predicted experimental effects, as well as make novel predictions about how prior learning biases action selection and affects reaction times. The key point will be to show that the sampling process is an integral part of the value-based decision making progress, i.e. observed reaction times provide a window into the inner workings of the planning agent. The agent evaluates behavior based on policies (action sequences) which allows us to not only illustrate behavior and reaction times in single trial experiments, but also learning of sequential behavior in a multi-trial sequential decision experiment. To present the model in a didactic fashion, we use a simple value-based decision making grid-world experiment [33, 34, 35].

For our simulations, the grid of the experiment consists of four rows by five columns yielding 20 grid cells (Fig 3A). Simulated agents start in the lower middle cell in position 3 (brown square) and have a simple task: to navigate to either one of the two goal positions at cells 15 (blue square) and 11 (green square) while learning their grid-world environment. Although the task would not be difficult for a human participant, this task gives us plenty of opportunity to illustrate how the model agent operates as a value-based decision maker and thereby generates choices and reaction times. In each cell, the agent has three options: move left, up and right. The tasks consists of 200 so-called miniblocks, where one miniblock consists of four trials. In each miniblock, the agent will start in cell 3 and is given the task to move to the indicated goal (either 1 or 2) within the four trials. During the first 100 miniblocks, goal 1 is active, and goal 2 is active in miniblocks 101 - 200. These two phases constitute two distinct contexts as they have different action–outcome contingencies. The agent is not informed about what cells give reward but has to find out by trial and error, inferring the contingencies of the current context.

**Fig 3.**
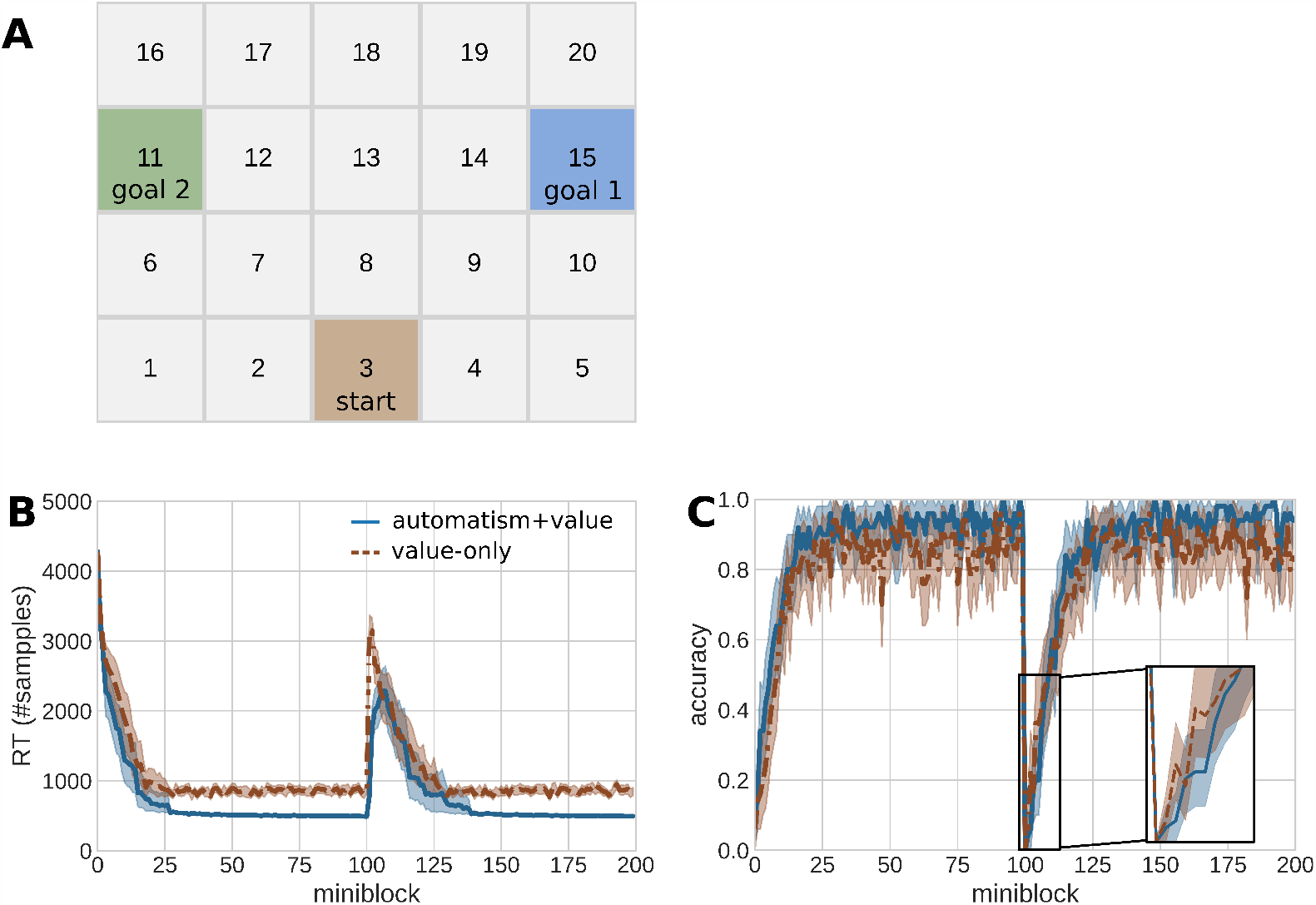
Reaction times and behavior in a sequential instrumental learning task. **A**: The grid world is an environment with 20 states. In each miniblock of four trials, the agent starts in the brown square (cell 3) and has to navigate to either goal 1 (blue square, cell 15) or goal 2 (green square, cell 11). The agent can use either of three actions in each step in the miniblock: left, up, and right. The experiment consist of 200 miniblocks. In the first 100 miniblocks, goal 1 is active, and in the second 100 miniblocks, goal 2 is active. **B**: Reaction times (as number of samples) of the first action in each miniblock over the course of the simulated experiment. The solid blue line shows the mean reaction time of 50 automatism+value learning agents (*a* = 1.0). The dashed brown line shows the mean reaction time of 50 value-only learning agents (*a* = 0.001). The shaded areas indicate a confidence interval of 95%. **C**: Accuracy in the same experiment, colors as in B. The accuracy was calculated as the percentage of miniblocks where agents successfully navigated to the currently active goal (goal 1 in trials 1-100, goal 2 in trials 101-200).

#### Reaction times and learning

As a first step, we show that simulated reaction times and choices generated by BCC agents behave as one would expect: In the beginning of the grid-world experiment, as well after a context change, RTs are high and accuracy (reaching the goal state) is low. They subsequently decrease and increase, respectively, as an agent familiarizes itself with the environment. Additionally, we show that, as expected, an agent with a higher automatization tendency has decreased reaction times and increased accuracy during the transition from goal-directed to more automatized behavior.

We divided 100 agents into 50 value-only learner agents (*a* = 0.001) (see Section Free parameters and simulation setups, and Supplementary 1: Detailed mathematical derivations) and 50 automatism+value learning agents (*a* = 1.0). The value-only learners adjust their prior over policies so slowly that the it plays effectively no role for action selection. In our simulations, we use the reaction time of the first action as an indicator how fast an agent decided to use a specific sequence of four actions to reach its goal. Fig 3B shows the mean reaction times per miniblock, for automatism+value and value-only learners. As expected, both agent types have slow reaction times in the beginning of learning how to reach the goal. This is because the agent does not know where the goal is yet, which means that all policies have equal value and hence sampling takes longer. The reaction times decrease strongly within the first 25 miniblocks, as agents become more certain about the goal location. Automatism+value learners additionally learn a pronounced prior over policies, thereby confining their action selection strongly, and displaying faster reaction times than the value-only learners. The reaction times of both agent types converge after around 50 trials to stable values, where value-only learners have generally larger and more variable reaction times.

Similarly, the mean accuracy (Fig. 3C) is initially low while agents learn about the environment, and increases strongly in the first 25 trials, until it stabilizes. The automatism+value learner agents achieve higher accuracy, as the stronger prior leads to a higher choice probability of the correct policy.

After trial 100, the contingencies of the environment change and goal 2 becomes active. Agents have to infer a new context and learn new goal-directed action sequences, as well as a new prior. This switching and learning of a new context is expressed in a large increase in reaction times and a drop in accuracy for both agent types, see Fig 3B&C. As in the first context, both agent types learn the new goal location within the first 25 trials, the mean reaction times decrease again to a stable value and accuracy also increases to a stable level. Here, the automatism+value learner again achieves lower reaction times and higher accuracy.

This shows that learning a strong prior over policies helps to achieve faster and more accurate behavior, as long as an agent is in a stable context. Only in the five trials after the switch (trials 101-105), it is disadvantageous to have learned a strong prior. The automatism+value learner has a decreased accuracy (0.096) in comparison to the value-only learner (0.164), see inlay in Fig 3C. This is because the agent assigns a non-zero posterior over the previous context after the switch, which leads to the old prior over policies interfering with choices in the automatism+value learner.

### Speed-accuracy trade-off and prior-based choices

In what follows we illustrate the internal sampling dynamics during the reaction time and choice generation in the grid world. We also show the influence of the speed-accuracy trade-off *s* and different constellations of prior and likelihood on choices and reaction times, see Section Planning and reaction times.

#### Sampling trajectories in the reaction time algorithm

Here we illustrate the sampling process of actions or policies in the proposed model. Fig 4 shows exemplary sampling trajectories. We show trajectories for 2 actions (top row) to illustrate how sampling works in more classical task settings, and for 81 policies from the grid world (bottom row, 81 = 3^4^ being the number of possible combinations of 3 actions that can be used in 4 time steps), using a smaller and larger speed-accuracy trade-off each. There are three interesting effects: First, the speed-accuracy trade-off *s* determines the time the estimated posterior has been close to the true value, before it commits to an action, see Section Free parameters and simulation setups. In the exemplary trajectories with a small trade-off *s* (Fig 4A&C), the reaction time algorithm commits rather quickly to a choice. In the trajectories with larger speed-accuracy trade-off *s* (Fig 4B&D), the sampling continues long enough for the estimated value to converge rather closely to the value of the true posterior before the sampling concludes.

**Fig 4.**
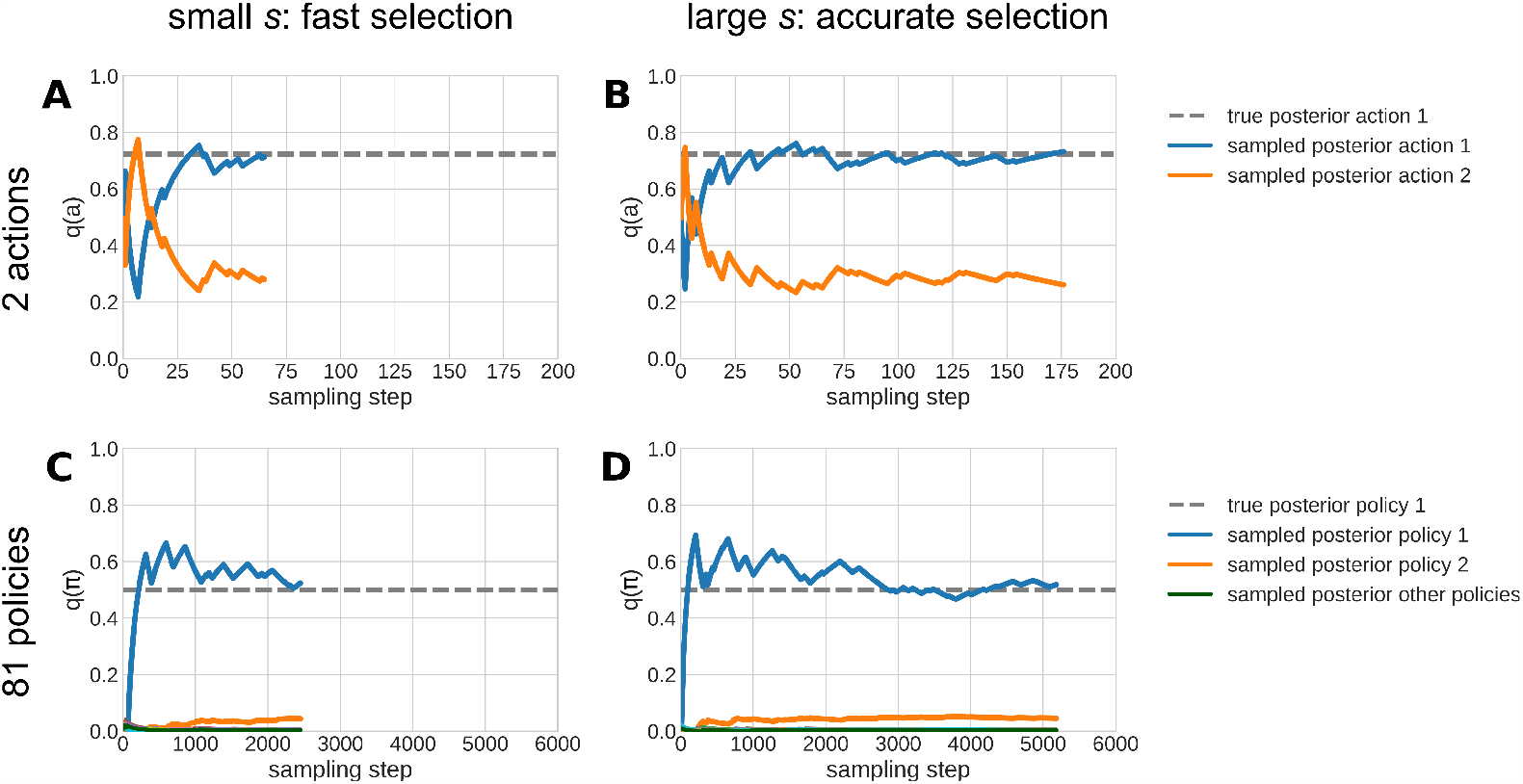
Reaction time sampling trajectories. **A** Exemplary sampling trajectories for a decision with two actions, with speed-accuracy trade-off *s* = 1.5. The dashed grey line shows the true posterior of action 1. The colored solid lines show values of the sampled posterior that emerge during the sampling in the reaction time algorithm. The blue line indicates the estimated value of action 1, and the orange line the sampled value of action 2. **B** Same as in (**A**) but with a larger speed-accuracy trade-off of *s* = 2.0 that results in more prolonged sampling duration. **C** Sampling trajectories for 81 policies (as is the case in the value-based decision task, see section Value-based decision making in a grid world), with a speed-accuracy trade-off of *s* = 0.5. Analogous to (A), the dashed line is the true posterior of policy 1, the blue and orange line are the sampled posteriors for policies 1 and 2, respectively, and the additional green line in the bottom row shows the estimated value of the other policies 3-81. **D** Same as in (C) but with a larger factor of *s* = 0.6. To illustrate the effect of priors, we set the prior of the second action (in A and B) or policy 2 (in C and D) to a greater value (0.6 for two actions, and 0.1 for 81 policies) than the prior of the first action or policy in all 4 panels (0.4 for two actions, and 0.9/81 for 81 policies). The likelihood and the resulting posterior always favor the first action and policy.

Second, one can see that the value of the estimated posterior fluctuates around the true posterior value (as for example in Fig 4A and B). This is because the sampling does not stop when the true posterior is crossed, but when the sampled value has been becoming closer for a number of sampling steps that are dependent on *s*. Due to the sampling being i.i.d draws of the true posterior, the estimated posterior converges towards the true posterior during the sampling process. During the sampling process, the estimated values may exhibit either oscillatory behavior, or an over- or undershooting followed by a convergence to the inferred value, respectively, see Fig 4C for an example with initial overshooting.

Third, for early sampling steps, the policy with the largest value in the prior may be sampled first, see Fig 4A&B. To illustrate this, we gave action 2 higher prior probabilities. That is why action 2 (orange line) is valued highly in the initial ten samples, despite having a lower posterior value than action 1 (blue line). For small speed-accuracy trade-offs *s*, this can lead to the model selecting a policy that has a large prior but a low posterior value. This is a setting that would reflect tight deadline regimes, e.g., when a response is required immediately, and thus automatic actions become more likely to be chosen. This is not always the case however, due to the inherent noise in the sampling. For example, despite being less likely, action/policy 1 may be chosen in the first sample due to the stochasticity of the sampling. This can be seen in the bottom row, where the sampling recognized policy 2 as better from the start, which leads to the estimated value of policy 2 simply being larger than the other policies in the bottom row.

In the effects shown here, one can see a key difference to an RRL+DDM approach. In the DDM, a choice is selected when the decision boundary is first crossed, whereas in the BCC model, a choice is made only when the estimated value has been sufficiently close for long enough. Importantly, the fluctuations in the trajectory of the proposed model are not simply added noise as in the DDM, but are an inherent part of the process of evaluating the value of each action. Each sample that moves the estimated value *p*(*π*|*ϑ*) up or down, corresponds to an evaluation of this policy in terms of prior bias and expected reward in the likelihood. In other words, the noise-like dynamics of the sampled trajectory, and consequently the resulting reaction times, are an essential part of the underlying action selection process.

#### Reaction time distributions and related choices

In this section, we show that the BCC is able to emulate classical log-normal reaction time distributions. We illustrate how the resulting distributions depend on the speed-accuracy trade-off *s* and the configurations of prior and likelihood. We furthermore show that shorter reaction times and smaller trade-offs *s* lead to more automatic choices from the posterior.

Fig 5 shows reaction time distributions for different trade-offs *s* = {0.1, 0.3, 0.5} given different combinations of prior and likelihood, as well as how similar samples are to prior and posterior, respectively. Overall, mean reaction times (in number of samples) are shorter and distributions are narrower for lower values of *s*. The distributions are either very narrow, or clearly right-skewed, the latter of which is typical for reaction time distributions. Indeed, all distributions pass a log-normal test with *p <* 0.01^2^. Mean RT and distribution variance furthermore depend on the configuration of the prior and likelihood that were given.

**Fig 5.**
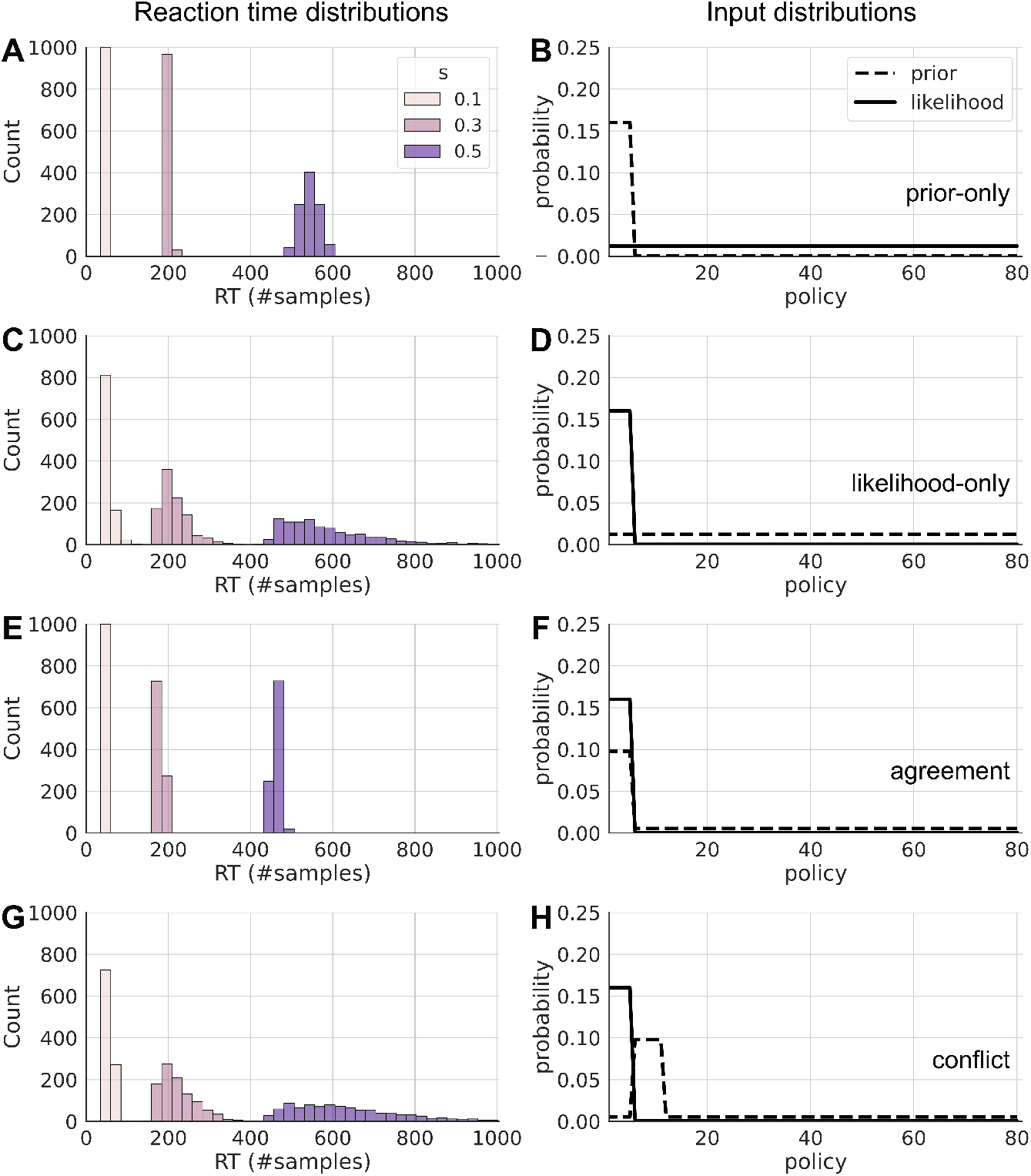
Reaction time distributions under different conditions. Reaction time distributions (left) for different input configurations of prior and likelihood (right), for different values of the speed-accuracy trade-off. **A**: Histogram of reaction times as number of samples in a prior-only, i.e. fully automatic setting, for *s* = {0.1, 0.3, 0.5} in beige, pink, and purple, respectively. The prior and likelihood that were used for the sampling are shown in B. The distributions have generally larger means and variance with increasing *s*. **B**: Probability values of the prior (dashed line) and the likelihood (solid line). The first six policies have increased prior values (taken from a grid world agent) while the likelihood is uniform. The posterior and resulting choices will be fully prior-based and therewith automatic. **C** Histogram of reaction times for a likelihood-only, i.e. fully goal-directed setting, input distributions are shown in D. Here, the distributions have a larger variance compared to A. **D**: Probability values of prior and likelihood. The likelihood is pronounced only for the six goal-leading policies in the grid world. The values of prior and likelihood were taken from the value-only learner in the grid world. **E**: Histogram of reaction time distributions for a setting, in which prior and likelihood are in agreement, as for example in the automatism+value learner in the grid world. Input shown in F. The resulting distributions have lower mean and variance than in the other settings. **F**: Probability values for prior and likelihood. Values were taken from the automatism+value learner in the grid world. **G**: Histogram of reaction times for a setting where prior and likelihood are in conflict (input values shown in H). This setting leads to increased mean and variance, as well as to skewed distributions. **H**: Prior and likelihood where the likelihood still favors the first 6 policies, while the prior biases towards other 6 policies. This results in a conflict, where goal-directed information and automatic biases are contradicting.

#### Goal-directed and automatic behavior

The first row (Fig. 7A and B) shows reaction time distributions for a prior and likelihood configuration where only the prior is pronounced while the likelihood is flat and uniform. This is a setting where automatic behavior from the prior will dominate choices. Vice versa, the second row (Fig. 7C and D) shows reaction time distributions for a configuration where only the likelihood is pronounced, while the prior is uniform. This is a setting where automatic biases play no role, and behavior will be dominated by goal-directed values. This configuration also equates to what a value-only learner naturally learns in the grid world above. Note, that for both, the resulting posterior, from which an agent samples actions, is the same, but the resulting reaction times for the likelihood-only case have much larger variance, but a similar mean. This shows that the proposed BCC model makes different predictions in terms of reaction time distributions, even if the choice probabilities are the same, showing the added value of taking reaction times into account.

The third row (Fig. 7E and F) shows a configuration, where prior and likelihood are in agreement. This is a configuration that naturally occurs for the automatism+value learner in the grid world. The resulting reaction time distributions (Fig. 7E) have the lowest mean and variance in the settings presented here. This is because the resulting posterior has very little uncertainty, and the samples from the prior agree with the values in the likelihood, so that the stopping criterion will be reached quickly.

#### Conflicts

A well-known phenomenon in reaction time experiments is that reaction times are longer in conflict trials, when one source of information is incongruent with another. For example, a conflict arises when the goal switches at trial 101, see Fig 3B, when the agent has learned a prior which is no longer useful for the changed goal location. The fourth row (Fig. 7G and H) illustrates such a conflict setting, where likelihood and prior point towards different policies. The resulting distributions have more variance and a higher mean compared to all other cases. This is because the posterior has more uncertainty, and the sampling will often draw policies from the prior, which have a low goal-directed value in the likelihood, leading to rejected samples. Conversely, it will take a large amount of sampling steps until a policy with a higher likelihood is sampled.

Fig 6 illustrates that the actions chosen by the agent in the conflict case are, for smaller speed-accuracy trade-offs *s*, more likely to be automatic, compared to larger *s*. The sampled policies up to the point of the choice in the chain have much more similarity to the prior for smaller *s*, i.e. a lower Kullback-Leibler divergence (DKL) between samples in the chain and the prior. For larger *s*, the samples in the chain resemble more the actual posterior. This shows that for tighter deadlines, or shorter reaction times, the chosen actions will be more automatic and less goal-directed.

**Fig 6.**
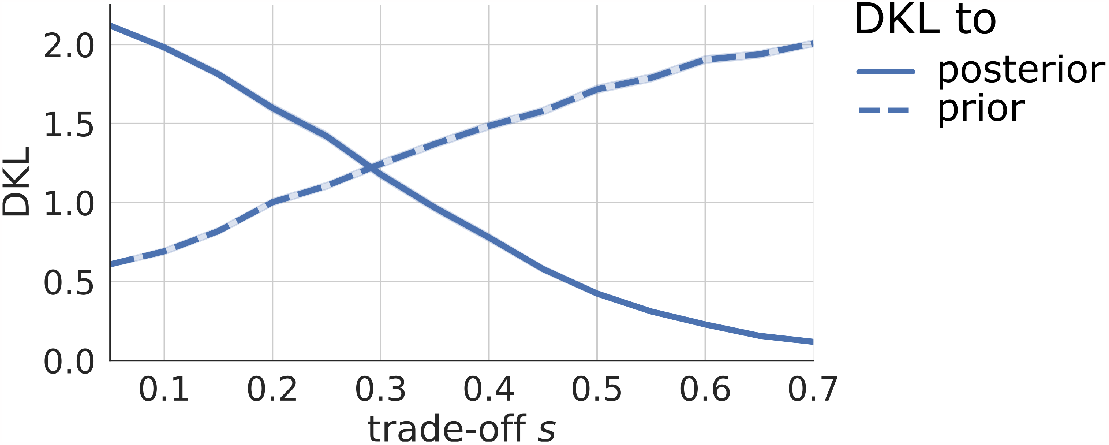
Similarity of samples with prior and posterior during conflict. The Kullback-Leibler divergence (DKL) between the samples in the chain and the true posterior (solid line), or the prior (dashed line), as a function of the speed-accuracy trade-off *s*, in the conflict setting from Fig 5H. The DKL between the samples in the chain and the prior decreases with *s*, whereas the DKL between the samples and the posterior increases with *s*. Hence, for lower values of *s*, the samples in the chain correspond more closely to the prior, which means that for shorter reaction times (e.g. the beige bars in Fig 5H) the chosen policy and corresponding action will be determined by the prior, and therewith automatic. For larger values of *s*, conversely, the samples in the chain resemble the true posterior more, so that for longer reaction times (e.g. the purple bars in Fig 5H), the sampling has converged and choices will be made in accordance to the posterior, taking goal-directed information into account.

### Flanker task

In this section, we show an application of the BCC model to a task that is well known to induce and measure conflicts: The Eriksen flanker task, which is a widely used behavioral task to measure inhibition under response conflicts [2]. Typically, in this task participants learn a stimulus-response mapping where one or two stimuli correspond to pressing one key, e.g. right, while one or two different stimuli correspond to pressing another key, e.g. left. The stimulus determining which answer is correct is typically shown in the middle of the screen. Conflict is introduced by showing distractor stimuli (flankers) surrounding the relevant stimulus in the middle. The distractors are chosen to be one of the stimuli that also indicate correct and incorrect responses. This induces congruent trials where the distractors indicate the same key as the relevant stimulus, and incongruent trials where the distractors indicate the other key.

It is typically found that RTs are increased while accuracy is decreased in incongruent trials, compared to congruent trials. The classical explanation of this effect is that, early in visual perception, all stimuli are processed in parallel, while perception focuses on the relevant stimulus in a later phase [4]. Here, we want to show that this effect can be understood in terms of the BCC model, where we do not model the perceptual process explicitly, but interpret flankers in terms of context cues and task stimuli in terms of goal-directed information. Specifically, we interpret the flanker distractor stimuli as context cues which indicate a “correct response” context, because the flankers and targets use the same stimuli, so that the flankers trigger an automatic response towards one key. To realize this in the model, we map policies to single actions, so that in each trial, an agent evaluates two policies each containing only one action. We implement an automatism bias through the automatization tendency in the prior over actions (see Section Free parameters and simulation setups, and Supplementary 1: Detailed mathematical derivations) to encode cue–response associations, and therewith implement direct flanker–response associations. In congruent trials, the prior and likelihood will be in agreement (see Fig 5E&F), which facilitates fast and accurate action selection. In incongruent trials, the prior is in conflict with the actual goal-directed stimulus encoded in the likelihood, which increases RTs and error rates, see Fig 5G&H. Importantly, to implement trial-wise learning, this inference setup is continuously updated throughout the experiment through a small forgetting factor (which is the Bayesian equivalent of a learning rate, see Free parameters and simulation setups and Supplementary 1: Detailed mathematical derivations). This leads to updating and re-learning of the flanker–response associations during the course of the experiment. In our simulations, we used a flanker task version with four compound stimuli. In the following, we replicate two well-established effects in flanker experiments.

#### Conditional accuracy function

One typical finding in flanker tasks is the conditional accuracy function (Fig 7A) [5]. In congruent trials, responses have a high accuracy independent of RT. However, in incongruent trials, responses with short RTs are most likely incorrect with the accuracy dropping below chance. There is even a linear relationship between RT and accuracy between 300 and 500ms, see the dashed line in Fig 7A. Fig 7B shows the averaged accuracy function of 50 simulated agents which shows a similar dip below chance level for low RTs in incongruent trials. While there are quantitative differences between experimental and simulated accuracy function, we can capture the key qualitative differences between congruent and incongruent trials. In the model, these effects are emulated because, for lower RTs, choices are more often made in accordance with the prior, while for long RTs, choices are mostly made in accordance with the posterior (see also Fig 5I).

**Fig 7.**
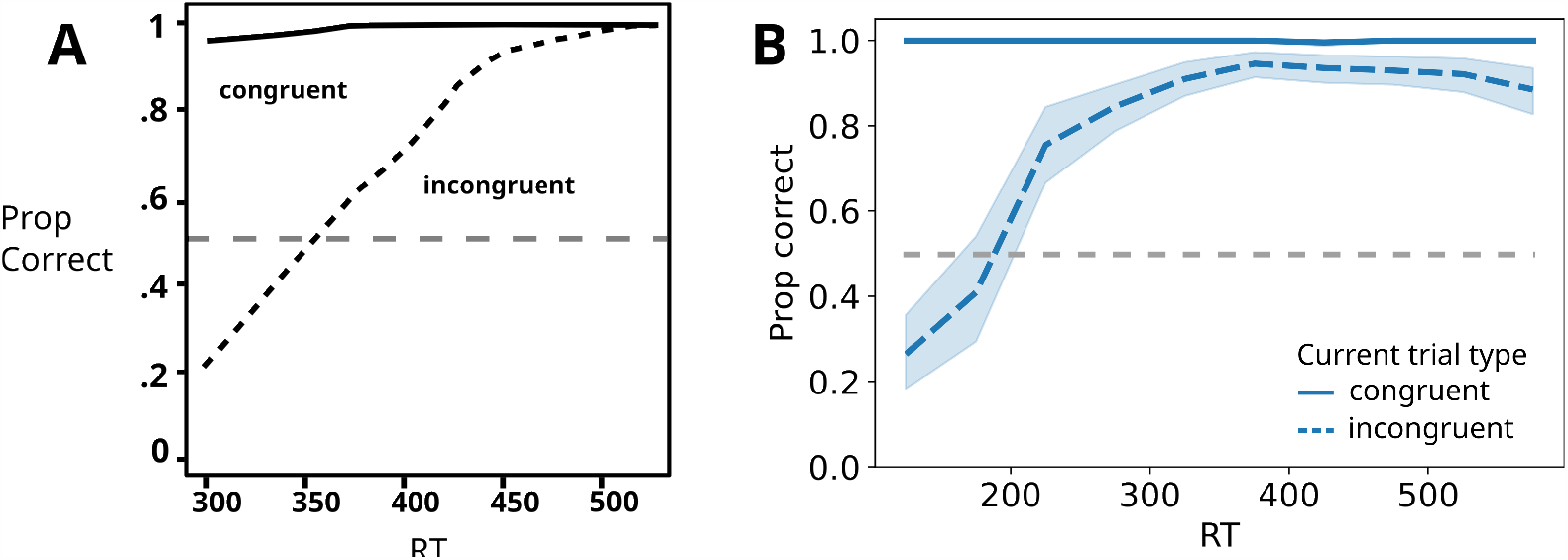
Conditional accuracy function. **A**: Conditional accuracy function in the flanker task, figure adapted from [36]. The solid and dashed lines show the proportion of correct responses for congruent and incongruent trials, respectively, as a function of reaction time. The gray dashed line indicates chance level. **B**: Simulated accuracy function, the line styles are as in A. The lines indicate the mean proportion of correct responses of 50 agents. The shaded area shows the confidence interval of 95%. Due to the congruent trial responses almost always being correct, the confidence interval is small and vanishes behind the solid line. The proportions were calculated by binning reaction times.

#### Gratton effect

Another typical finding in the flanker task is the so-called Gratton effect (Fig 8A) [4]. Here, mean reaction times decrease in the second consecutive trial of the same type (congruent vs incongruent). Usually, the Gratton effect is interpreted as being due to different degrees of recruitment of cognitive control depending on the previous trial type [2]. In the BCC, we explain the Gratton effect as a sequential effect, which is due to strengthened or weakened associations between the distractor (context) and the response. A congruent trial strengthens this association, making prior and likelihood more or less similar in a following congruent or incongruent trial, respectively. This in turn decreases or increases reaction times respectively. In Fig 8B, we show that we can replicate the Gratton effect using this mechanism.

**Fig 8.**
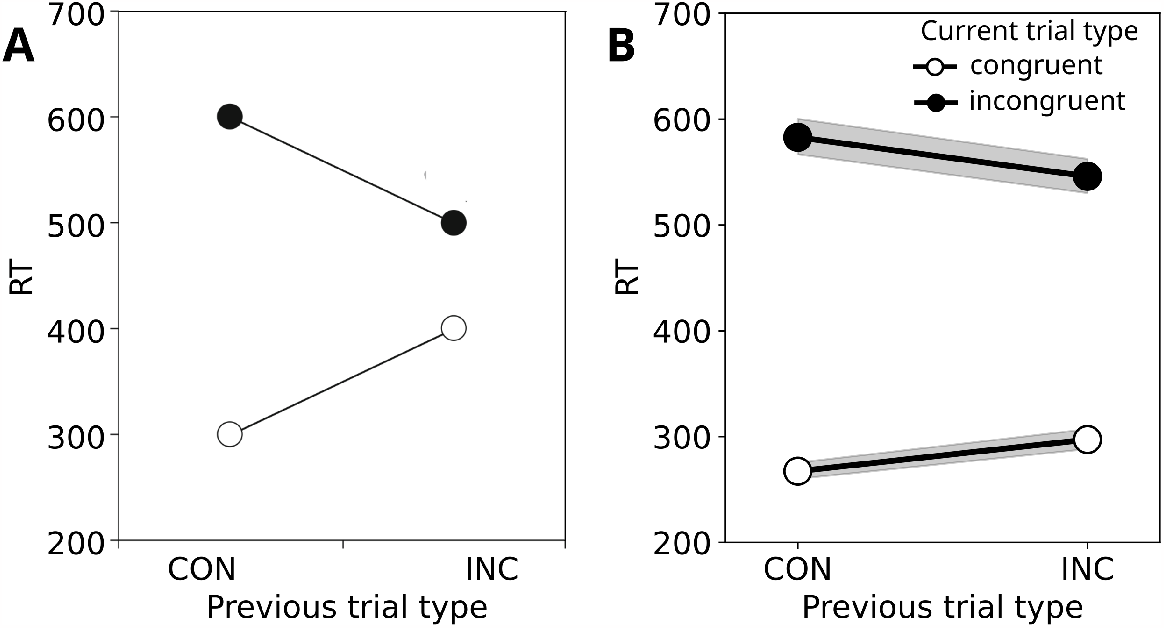
Gratton effect. **A**: Typical Gratton effect findings in the flanker task, figure adapted from [37]. The x-axis shows the previous trial type, either congruent (CON) or incongruent (INC). The circles indicate the current trial as either incongruent (black) or congruent (white). Reaction times on the y-axis differ between congruent and incongruent trials, and depending on the previous trial type. **B**: Simulated Gratton effect. The axes and colors are as in A. The shaded areas show a confidence interval of 95%. Note that we use here Eq 9 to convert number of samples to milliseconds, so that RT is in ms as in A.

In order to demonstrate that the setup of the prior, the experience-dependent updating of the prior, and importantly, its interplay with the context are the key machinery which allows us to simulate typical flanker effects with the BCC model, we show in the Supplementary material Supplementary 2: Flanker features that leaving out any of these three components will drastically change the conditional accuracy function and nullify the Gratton effect.

### Task switching

Besides the conflict effects shown above, contexts can interfere with each other in other ways, for example by experiencing a succession of different contexts, where it is required to switch and load different context properties. This is the domain of so-called task switching tasks, which measure task set shifting and switching. In a typical task switching task [6, 7], participants are presented with two different response rules. For example participants must indicate whether a number is even or odd, and whether a letter is a vowel or a consonant. For a trial, the current task set is cued, and a stimulus is presented with features of both task sets, such as a letter and a number. The participant has to respond to the stimulus under the current task set. Due to the stimulus consisting of both features, trials can be congruent or incongruent: In congruent trials, both features require the same response, and in incongruent trials the two features require different responses. In this task, participants experience two sources of uncertainty: (i) The task (context) cue may be perceived or processed in a noisy manner, which is also influenced by the cue presentation time, i.e. how long the task cue is visible before a response is warranted, and (ii) uncertainty about the upcoming task, and how often the task changes.

In the BCC model, task switching corresponds to switching between two different contexts with different outcome rules. The agent learns the outcome rules for each context in the likelihood at the beginning of the experiment and later in the experiment loads the task-specific learned rules in response to the task set (context) cue. The agent is set up such that context cues are observed with low but non-zero uncertainty emulating a limited cue presentation time and perceptual uncertainties. The agent’s expectation of context transition probability is set to be relatively low as well, to a regime in which context inference is stable but the recognition of a context switch is still possible (see also Section Free parameters and simulation setups). The resulting low but existing uncertainty about the current context results in traces of the previous context still being present in a switch trial. These traces fade with each consecutive trial in the same context. The previous context may therefore introduce conflicts which increase reaction times, as is typically observed in this task. Note that here, we use a value-only learner (*a* = 0.001) to focus less on learning a prior, just as a human participant would probably do when there is no discernible repetition of one choice over another.

In this section, we replicate three common findings from the task switching literature [6, 7, 38, 39]: (1) decreased reaction times in repetition trials of the same task set, (2) decreased reaction times with longer cue–target intervals, and (3) decreased reaction times with longer response–stimulus intervals.

#### Repetition trials

In the first trial after a task switch, reaction times typically increase. This finding is typically interpreted as being caused by underlying costs associated with switching the task set. In particular, the previous task set may interfere with the response to the new task set. Additionally, as in the Flanker task, participants are typically slower in incongruent trials than in congruent trials. Lastly, reaction times typically decrease as more trials in the task have been trained. These results are shown in Fig 9A. Fig 9B shows simulated average reaction times of 50 agents over the course of a 70 trial task switching experiment. As in the experimental findings on the left, shorter periods of training lead to larger reaction times. While simulated reaction times are noisier and do not quite have the same absolute values, incongruent trials lead to increased reaction times, especially in switch trials both, in the experiment and the model. In addition, reaction times decrease as a function of repeated trials.

**Fig 9.**
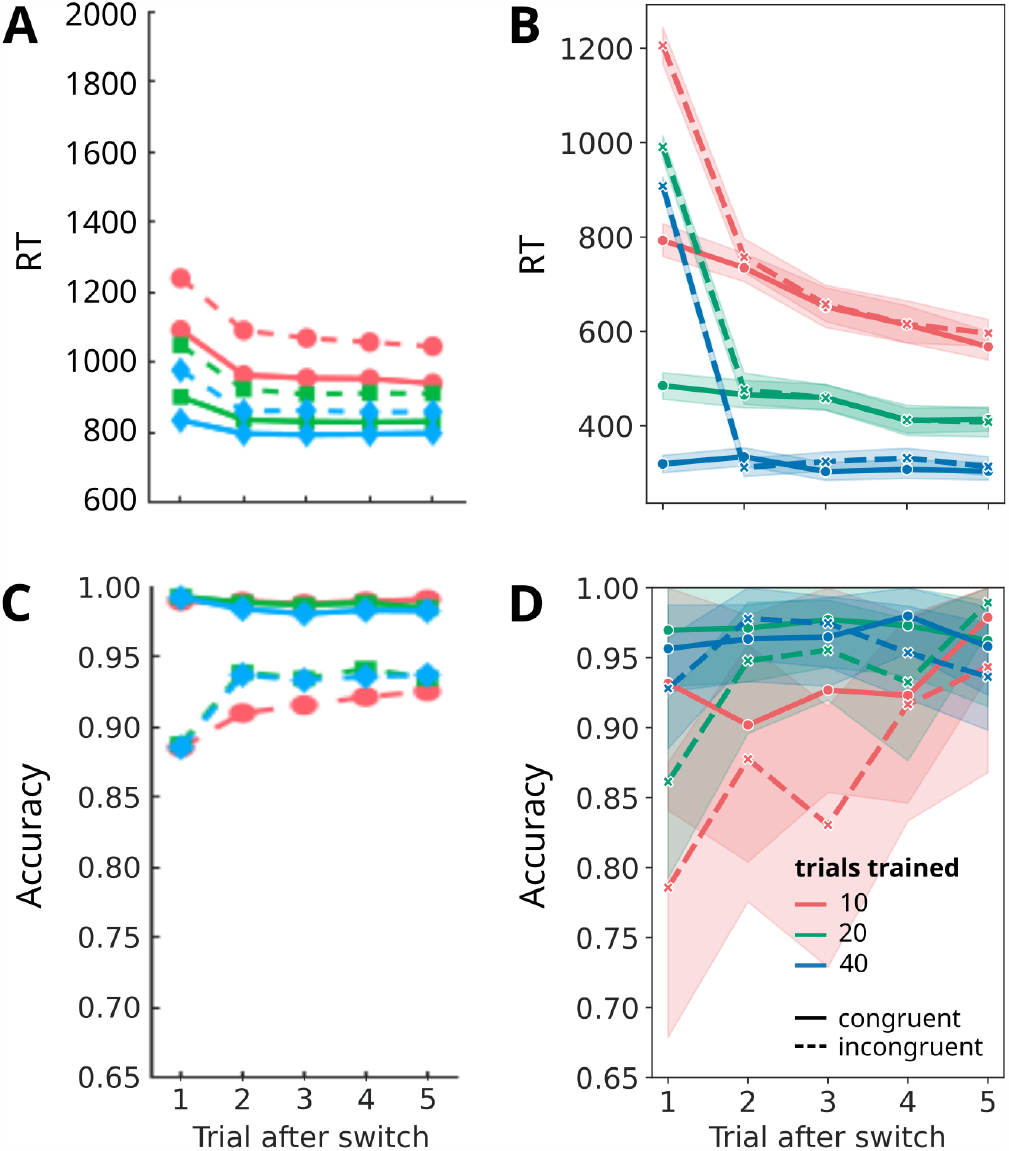
RTs and accuracy as a function of repeating trial after switch. **A**: Reaction time in a typical task switching experiment as a function of trial number after the switch, adapted from [38]. Red lines indicate little previous training (*∼*10 trials), green lines indicate medium training (*∼*20 trials), and blue lines long training (*∼*40 trials). Dashed lines are incongruent trials, and solid lines congruent trials. The shaded areas correspond to a 95% confidence interval. **B**: Simulated average reaction times from 50 agents. Colors as an A. **C**: Response accuracy as a function of trial number after switch from the same experiment as A. Colors as in A. **D**: Simulated average response accuracy from the same simulated experiment as in C. Colors as in A.

In the agent, these effects are due to the previous task, i.e. context, lingering because of the remaining uncertainty about contexts. The longer a task was active, the lower the remaining uncertainty, and the less the outcome contingencies of the previous task influence the decision. This effect is stronger in conflict trials, as the contingencies of the other task may contradict the contingencies of the current task. This induces uncertainty in the action selection and therewith increases reaction times.

The converse has been found experimentally for response accuracy: Accuracy drops in switch trials, and is lower in incongruent trials (Fig 9C). In Fig 9D, we show the simulated average accuracy. The accuracy difference between congruent and incongruent trials is not quite as large in the simulated data compared to the experimental data, but is clearly present for the 20 and 40 trials trained case (green and blue lines) and is mostly present for the 10 trials trained case (red line).

To demonstrate that context inference and context–dependent learning are essential for modeling a task switching task, we show in Supplementary 3: Task switching features that removing the context feature leads to vastly different reaction time and accuracy effects compared to those shown here, which do not resemble the typical task effects anymore.

#### Cue–target interval

Another well known effect in task switching is the influence of the cue–target interval (CTI) on reaction times [39]. The cue–target interval is the time between the presentation of the current task cue and the onset of the stimulus (target) after which a response has to be made. Longer CTIs allow participants to better process the cue and load the task set before the onset of the stimulus. Fig 10A shows how reaction times increase with shorter CTI, compared to a single task experiment [39].

**Fig 10.**
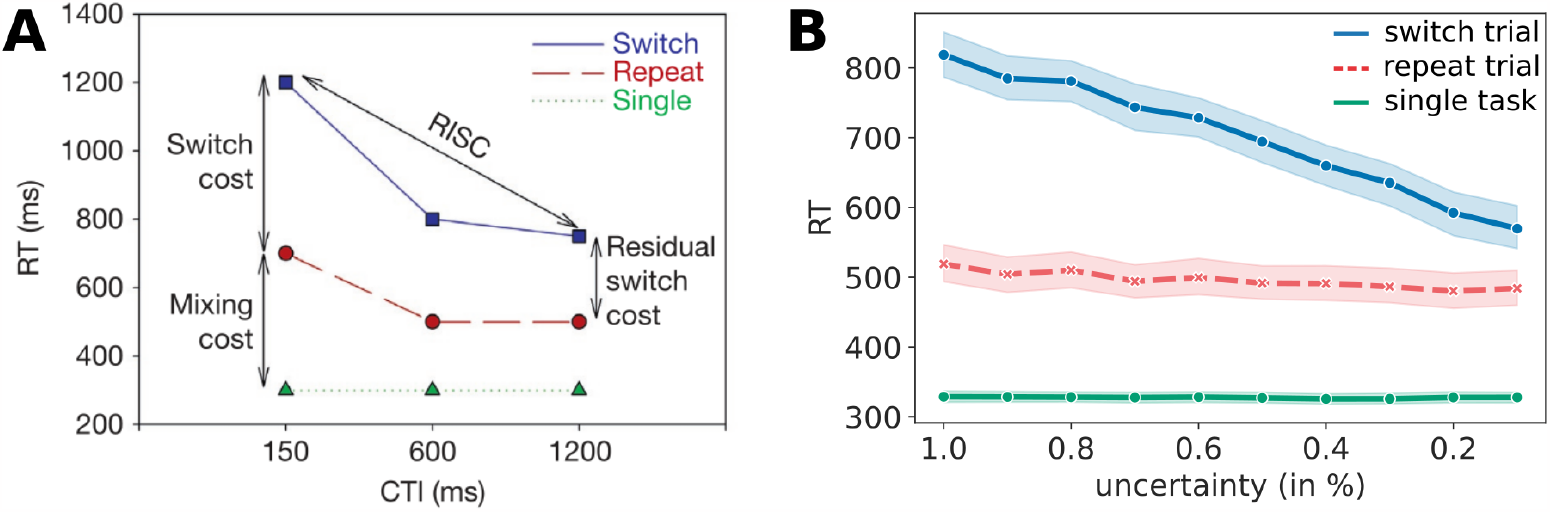
Cue–target interval. **A**: Reaction times in a single task experiment (green), in repeat trials in a task switching experiment (red), and in switch trials (blue), as a function of CTI; adapted from [39]. **B**: Mean simulated reaction times from 50 agents, as a function of cue uncertainty, colors as in A. The shaded ares indicate the 95% confidence interval. Note that for the green line, the confidence interval is narrow and is obscured by the line itself.

To model the CTI, we map shorter CTIs to higher uncertainty when perceiving the context cue that indicates the current task (see Section Free parameters and simulation setups, and Supplementary 1: Detailed mathematical derivations). Fig 10B shows average reaction times in switch trials, repeat trials, and trials in a single task experiment as a function of context cue uncertainty. Using this setup, we were able to qualitatively replicate the typical shape of CTI effects. The higher uncertainty during perception and processing of the task cue leads to increased reaction times because the previous task’s contingencies have higher influence on action selection.

#### Response–stimulus interval

A similar effect has been observed when varying the response–stimulus interval (RSI) [7], which is the time between a response and the next trial. Longer RSIs allow participants to better prepare for a possible switch and as a result reaction times decrease. Fig 11A shows experimentally established reaction times in switch and repeat trials as a function of the RSI [7].

**Fig 11.**
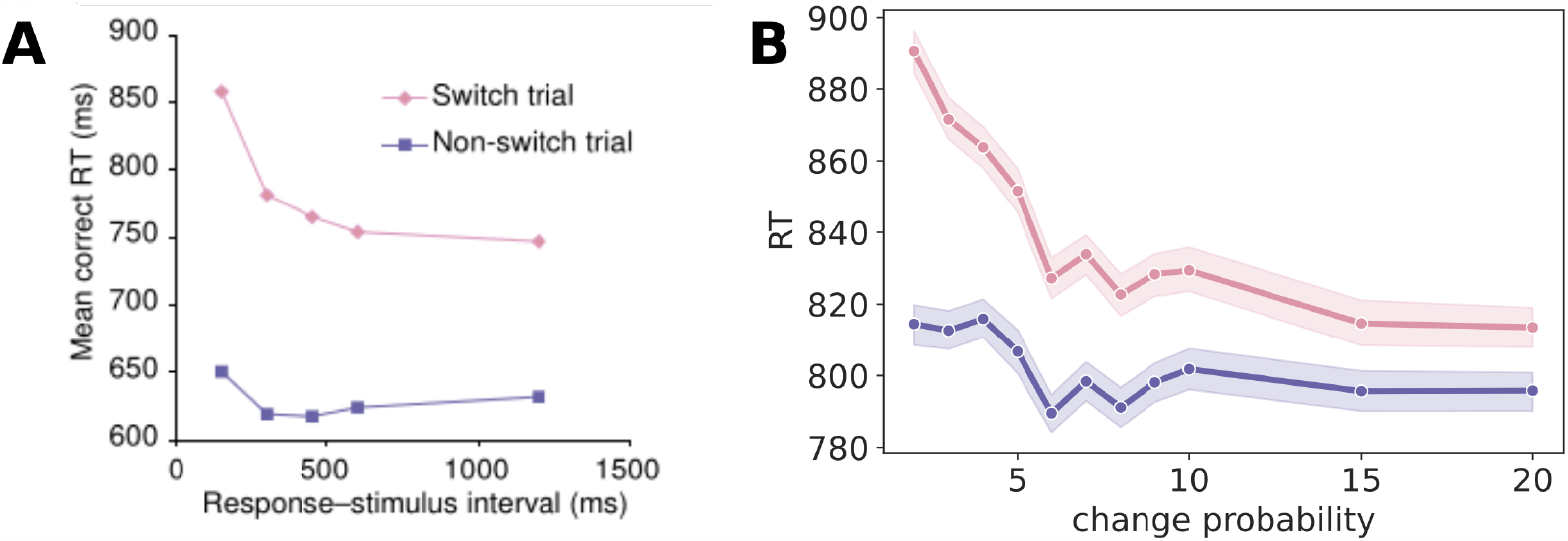
Response–stimulus interval. **A**: reaction times as a function of RSI for switch trials (pink) and repeat trials (purple), taken from [7]. **B**: Mean simulated reaction times from 50 agents, as a function of change probability, colors as in A. The shaded areas indicate the 95% confidence interval.

We model RSI differences as different levels of an agent’s assumption about context change probability in between trials (see Section Free parameters and simulation setups, and Supplementary 1: Detailed mathematical derivations). The higher the change probability, the easier it is for an agent to infer that the context and task changed, and to load the new task set and respond accordingly. Fig 11B shows average reaction times as a function of context change probability. Although there are quantitative differences, a similar pattern of decreasing reaction times as in the RSI emerges.

## Discussion

We have proposed a novel joint behavioral model, the Bayesian contextual control (BCC) model, that describes both choices and reaction times as measured in behavioral experiments. Planning corresponds to estimating posterior beliefs over policies (sequences of actions) using a posterior sampling scheme, Bayesian independent Markov chain Monte Carlo (MCMC). Posterior beliefs over policies are proportional to policy likelihood (a function of rewards) and an adaptive policy prior that tracks past actions [24]. MCMC sampling of the posterior beliefs terminates in accordance with the precision of posterior beliefs, and the resulting reaction time is computed from the number of samples. We first presented results in a grid world toy experiment in which we showed basic expected characteristics of this model and illustrated three key properties of the resulting reaction times: (1) Decreases in reaction times while learning a sequential value-based decision task; (2) shorter reaction times when automatic behavior is learned and when speed-accuracy trade-off is low; (3) classic log-normal shapes of reaction times, and how they relate to internal features of the decision process. We also showed that the BCC can qualitatively replicate findings in two cognitive control experiments: For a flanker task, we emulated the reaction time dependent conditional accuracy function, as well as the sequential Gratton effect. For a typical task switching task, we replicated repetition effects, cue–target interval (CTI) effects, and inter-trial (response–stimulus interval, RSI) effects.

In the BCC model the environmental context determines the current set of state transition and outcome rules. The rules are specific to the current task set, and are used to compute expected rewards in the policy likelihood, which encodes goal-directed information. We found context to be crucial for modeling the dynamical structure of task switching (see also Task switching). As the context itself is a discrete latent variable, the agent forms beliefs over possible contexts and assigns precision to those beliefs. Imprecise beliefs over contexts are critical for observing behavioral effects in the task switching paradigms, as both the effects of the cue–target interval as well as the response–stimulus interval depend on context inference uncertainty.

Another important aspect of the BCC model is the learning of context-specific implicit biases, in the form of priors over policies. These contextual policy priors are independent of goal-representations and follow the statistics of past choices. Contextual policy priors lead to automatic behavior which makes choices both faster and more reliable (see Value-based decision making in a grid world). Additionally, associating flankers with implicit context-response biases (where cue sets a context) results in recognizable response patterns typically observed in the flanker task, see Flanker task. Implementing automatic behavior as a context-dependent prior over actions or policies allows for repetition-based learning of context–response associations [40].

Combining these aspects of the BCC model with a sampling-based planning and action-selection, we were able to generate reaction times and and represent reaction-time distributions. During sampling, the prior provides a quickly available heuristic for estimating which parts of the decision tree should be evaluated first, and which can be ignored; effectively implementing priority-based sampling. This way, the prior not only encodes context–response associations but also helps to confine the space of what behavior to evaluate in a goal-directed manner. If the prior probability of a specific policy is close to zero, as would in a real-world setting be the case for many policies, these are effectively a priori excluded and will most likely never be sampled nor evaluated. The prior therewith helps to save computational resources and time when sampling the likelihood of actions. This mechanism sheds light on how learning speeds up decision making and saves resources by biasing the planning process itself.

Another advantage of this mechanism is that, during sampling, at every step a decision can be made, simply by executing the currently sampled policy as a heuristic. This would allow for fast but accurate action selection even in settings with unexpectedly short deadlines, such as typical situations in traffic. Policies sampled early in the process would typically be an adequately good choice to execute, due to the prior-based sampling.

### Implications and predictions

Although typical cognitive control tasks are usually not interpreted to rely on value-based decision making, the proposed modeling approach leads to an interesting view on cognitive control, which is typically interpreted as top-down control [41, 42, 1, 2]. In contrast, in the model, bottom-up inference of the current context plays an important role as well. Here, an inferred context determines not only which task rules currently hold but also which policies should be preferred in the current situation. Consequently, we model contexts at a higher level in the hierarchy, relative to actions. This enables the agent to probabilistically infer the high-level state context from its sensory inputs. In this sense, cognitive control, from a modeling perspective, is not only about control but also about inference [25, 26]. In the case of a high posterior on a specific context, ‘cognitive inference’ determines with high precision what rules the agent should currently follow, and which a priori information to use [43]. This behavior would be interpreted by an outside observer as highly controlled. In the case of a low posterior, with some weight on other contexts, behavior may look not too well-controlled, simply because the person cannot infer the current context well. In other words, precisely inferred contexts may look to an observer like ‘top-down control’ but may be understood as the agent’s recognition of the current situation as a function of context inference and having learned how to behave in it, given previous exposures to the same or similar situations [44].

Besides these theoretical implications, there are some interesting predictions about how traits that are quantified in this model would influence information processing in the brain and consequently, learning and decision making, which could be tested experimentally. With the BCC, one cannot only model classical experiments as shown here, but also in principle model choices, reaction times and learning trajectories in multi-step [8, 10] and multi-choice experiments [45]. Using the differences in resulting reaction time distributions for different settings of prior and likelihood (Fig 5), as well as the increased mean reaction times due to internal conflict, the BCC could be used for inferring trait-like quantities from behavioral and reaction time data: How quickly and strongly a participant learns a prior, how well reward probabilities and action–outcome contingencies have been learned at any point in time, and how difficult or easy it is for a participant to switch to a new context. These trait-like quantities may be linked to maladaptive behaviors and disorders, which can be tested in future experiments.

### Relation to other joint behavioral models

The classical modeling approach for reaction times is the influential drift diffusion model (DDM) [17, 46, 32, 47] which is in cognitive science one of the most established textbook models [18, Chapter 3]. It has been successfully applied across many experimental domains, most notably to perceptual decision making. DDMs fall under the umbrella of evidence accumulation models which describe the process of action selection and resulting reaction times as a biased random walk with a drift and white noise [32, 47] and have recently been extended for multi-choice tasks [14, 16]. As described above (see Fig. 2), this model type has been recently combined with reinforcement learning to provide joint value-based and reaction time models, where typically internal quantities of value-based model are linked to variables in the DDM (for example Q-values and drift rate) [11, 12, 13, 14, 16], see also Fig. 2. Learned values or value differences drive the random walk until a boundary is reached and the respective choice is executed.

However, the two models in this case are rather two separate mechanisms which are linked in a feed-forward way, and not by an integrative sampling mechanism as in the BCC. Additionally, the DDM does not provide an internal mechanistic explanation for the white noise component, which is simply added in each step. As it stands, this noise component in DDMs is a useful modeling device to explain reaction time distributions but it is typically not linked to an underlying generative mechanism within the model.

Consequently, a key difference of the DDM+RL-based approach to our proposed model is that we use sampling to not only provide a way to generate reaction times from the choice values of the value-based model, but to describe a potential mechanism how the inherent noise of the sampling contributes to the actual decision process, of which reaction times are a measurable by-product.

An important point is that sampling in the proposed model continues until the agent is sufficiently certain that the sampling has yielded enough information on what outcomes to expect for behavioral policies, see Section Sampling trajectories in the reaction time algorithm. The level of certainty is regulated by the speed-accuracy trade-off parameter *s*. This approach automatically integrates the uncertainty about outcomes into action selection, and is related to optimal stopping problems [48]. This makes the sampling qualitatively different from the sampling in typical evidence accumulation models, where sampling continues until a boundary is reached [17, 46, 32, 47]. In combined DDM+RL models, confidence and speed-accuracy trade off are implemented via time-varying decision boundaries [49, 12, 50, 51], which are in fact needed to ensure choice optimality for the combined model [12, 50, 51].

Additionally, the Bayesian nature of the BCC model leads to reaction times being a reflection of how similar or dissimilar the prior and the likelihood are in a specific context, and longer reaction times mean that a decision maker took longer to find a compromise between these two influences. Other simulation studies which used Bayesian or active inference models have also related reaction times to the dissimilarity of distributions [22, 23]. Contrary to the BCC model, these studies related reaction times to the dissimilarity of prior and posterior. This approach is closely related to our approach, as the likelihood ultimately determines the posterior, given a prior. Note that in both studies [22, 23] the authors did not propose a concrete mechanism how reaction time distributions are generated from prior and posterior, but focused on the principle of this relation. Hence, the BCC could be regarded as an extension and practical example of this approach, using an MCMC sampling mechanism.

MCMC was chosen as a sampling algorithm because it is a well-established general method to sample from probability distributions. Additionally, it has been argued recently that probabilistic computations may be implemented by neurons in the brain via types of MCMC sampling [52, 53, 54]. In line with this view, we interpret the relatively large number of samples required to reach a decision in terms of neuronal sampling.

In this article, we focused on describing reaction time variability as caused by sampling in the decision and action selection process. However, it is also possible that additional RT variability may be generated by sampling in other domains, such as in the planning process itself which can be implemented via sampling [55, 56, 21], and perceptual processes [57, 58]. For example in the flanker task, past modeling approaches have focused on perceptual uncertainty to explain reaction time effects, e.g. [36] while we focused solely on the decision process itself. It is hence possible that explicitly modeling perceptual effects in the flanker task may further improve the match to observed reaction time distributions and effects.

## Conclusion

In this paper we proposed a novel integrative reaction time and planning mechanism based on Bayesian sampling. Using this approach, we showed that we can qualitatively replicate a range of different well-known behavioral effects. The model components, the prior, context, and sampling-based planning, have been infrequently used in the literature. Given our results, we hypothesize that the combination of these components are a promising basis for future models of behavior, or may even map to more general cognitive processes underlying planning and decision making.

## Supporting information

Supplement 1, mathematical derivations

Supplement 2, flanker details

Supplement 3, task switching details

## Supporting information

The code has been made publicly available on github: https://github.com/SSchwoebel/BalancingControl

**Supplementary 1: Detailed mathematical derivations**. We show how the equations the agent is based on are derived from a probabilistic model using variational inference.

**Supplementary 2: Flanker features**. We show that the context and learning of the prior are fundamental features of the model to generate flanker task effects.

**Supplementary 3: Task switching features**. We show that the context is a fundamental part of the model to generate task switching effects.

## Acknowledgments

We thank Maarten Jung for his help with a previous version of the MCMC sampling algorithm as part of his masters thesis (https://nbn-resolving.org/urn:nbn:de:bsz:14-qucosa2-740483).

## Funding acknowledgments

Funded by the German Research Foundation (DFG, Deutsche Forschungsgemeinschaft), SFB 940 - Project number 178833530, and TRR 265 - Project number 402170461, and by the Saxon State Ministry of Science, Culture and Tourism (SMWK) as part of support for profile-defining structural units of the TUD in 2023/24, and as part of Germany’s Excellence Strategy – EXC 2050 – Project number 390696704 – Cluster of Excellence “Centre for Tactile Internet with Human-in-the-Loop” (CeTI) of Technische Universität Dresden.

## Footnotes

1 https://github.com/SSchwoebel/BalancingControl

2 Log-normality was tested using the python scipy normaltest function on the log of the RTs.

