## Supplement 1, mathematical derivations for "Joint modeling of choices and reaction times based on Bayesian contextual behavioral control"

### Supplementary material 1: Detailed derivations

In the prior-based contextual control model, first introduced in (Schwöbel, Marković, Smolka, & Kiebel, 2021), an agent maintains a probabilistic generative model of the dynamics and causal relationships in its environment. In order to form beliefs unobservable variables, the agent inverts this model using variational inference. Note, that in this model, not only environmental variables, such as the context, are treated as hidden variables, but also the actions or action sequences (policies) that an agent may choose. In what follows we will outline the details of the generative model, how an agent forms beliefs using variational inference, and what factors influence action selection.

#### Generative model

The agent summarizes environment dynamics into two levels of a hierarchy: On the top level, there are partially observable contexts, which change on a slower time scale, in between behavioral episodes of an agent. On the lower level, each context maps to a specific Markov decision process (MDP), that an agent navigates by selecting actions. This MDP is of finite horizon $K$, constituting episodes of length $K$. In an episode, an agent can select $K-1$ actions, which can be summarized as deterministic policies $\pi$ of length $K-1$. Additionally, not all parts of the context-specific MDPs are predetermined. An agent learns the reward rules for a context, as well as a prior over policies for each context. See Figure 1 for a graphical representation of the generative model.

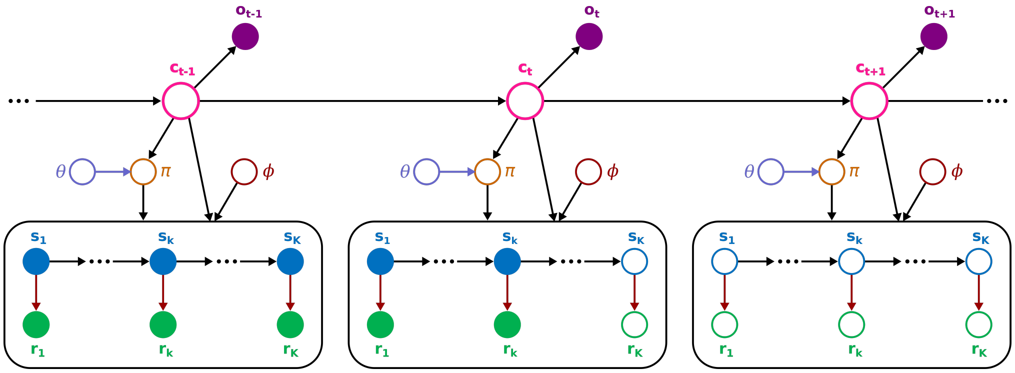


**Fig 1. Generative model** A graphical representation of the probabilistic generative model. Filled circles indicate observed variables, and empty circles indicate hidden variables. Arrows indicate conditional dependencies, and colored arrows show dependencies which are updated and learned. The lower boxes show MDPs of length $K$. The blue circles are the states of the MDP through which the agent navigates. States up until $\mathbf{s}_k$ of the current episode $t$ have been observed, while states from $\mathbf{s}_{k+1}$ have not yet been observed and are therewith hidden variables. Blue circles show rewards, which have been observed up until the current reward $\mathbf{r}_k$ in the episode $t$. The red arrows between the states and rewards indicate that the reward generation rules are parameterized by the set of variables $\phi$ (red circles), and these parameters are learned during an experiment. The policy $\pi$ (brown circle) is inferred executed by an agent, which determines the state transitions in the episode. The prior over policies is parameterized by a set of variables $\theta$, which are also learned during an experiment. On the upper level there are contexts $\mathbf{c}$ (pink circles) which determine the currently applicable outcome rules as well as the prior over policies. The context elicits cues $\mathbf{o}$ (purple filled circles).

The generative model can be written as

$$p\left(\mathbf{r}_{1:K}, \mathbf{s}_{1:K}, \phi, \pi, \theta, \mathbf{c}_t, \mathbf{o}_t\right) = \tag{1}$$

$$p\left(\mathbf{r}_{1:K}, \mathbf{s}_{1:K} | \pi, \phi, \mathbf{c}_t\right) p\left(\phi\right) p\left(\pi | \theta, \mathbf{c}_t\right) p\left(\theta\right)^{\lambda} p\left(\mathbf{o}_t | \mathbf{c}_t\right) p\left(\mathbf{c}_t\right) \tag{2}$$

where

$$p\left(\mathbf{r}_{1:K}, \mathbf{s}_{1:K} | \pi, \phi, \mathbf{c}_t\right) = \prod_k p\left(\mathbf{r}_k | \mathbf{s}_k, \phi, \mathbf{c}_t\right) p\left(\mathbf{s}_k | \mathbf{s}_{k-1}, \pi, \mathbf{c}_t\right) p\left(R | \mathbf{r}_k\right) \tag{3}$$

$$p\left(\mathbf{r}_k | \mathbf{s}_k, \phi, \mathbf{c}_t\right) = \prod_{r,s,c} \phi_{rsc}^{\delta_{r,\mathbf{r}_k} \delta_{s,\mathbf{s}_k} \delta_{c,\mathbf{c}_t}} \tag{4}$$

$$p\left(\mathbf{s}_k | \mathbf{s}_{k-1}, \pi, \mathbf{c}_t\right) = \prod_{s',s,c,j} p_{s'scj}^{\delta_{s',\mathbf{s}_k} \delta_{s,\mathbf{s}_{k-1}} \delta_{c,\mathbf{c}_t} \delta_{j,\pi}} \tag{5}$$

encodes the transition rules of categorically distributed states $\mathbf{s}_k$ and rewards $\mathbf{r}_k$, which depend on the categorically distributed current context $\mathbf{c}_t$ and the learned parameters $\phi$, which follow the corresponding conjugate Dirichlet prior

$$p\left(\phi\right) = \frac{1}{B\left(\beta^{t-1}\right)} \prod_{r,s,c} \phi_{rsc}^{\beta_{rsc}^{t-1} - 1}. \tag{6}$$

The concentration parameters $\beta^{t-1}$ keep track how often a reward $r$ has been received in a state $s$ when being in a context $c$ up until the previous episode $t-1$. Reward magnitudes are encoded in the dummy variable $p\left(R | \mathbf{r}_k\right) \in [0,1]$ which encodes how likeable a specific reward is. Additionally, state transitions are dependent on the chosen policy $\pi$. The policy $\pi$ has its own context-dependent categorical prior

$$p\left(\pi | \theta, \mathbf{c}_t\right) = \prod_{j,c} \theta_{jc}^{\delta_{j,\pi} \delta_{c,\mathbf{c}_t}}, \tag{7}$$

which depends on parameters $\theta$ that are Dirichlet distributed

$$p\left(\theta\right) = \frac{1}{B\left(\alpha^{t-1}\right)} \prod_{j,c} \theta_{jc}^{\alpha_{jc}^{t-1} - 1}. \tag{8}$$

The concentration parameters $\alpha^{t-1}$ keep track of how often a policy has been chosen in a specific context up until the previous episode $t-1$. Notice that $p\left(\theta\right)$ is exponentiated with a forgetting factor $\lambda \in [0,1]$ in the generative model (Eq. 2). This corresponds to assuming $\theta$ is a mixture distribution of learned concentration parameters $\alpha^{t-1}$ and a flat Dirichlet distribution, which leads to decreasing $\alpha^{t-1}$ over time, i.e. forgetting, for $\lambda < 1$. For a motivation and full derivation see (Moens & Zénon, 2019).

These variables therewith encode the MDP on the lower level of the hierarchy. Note that the prior over policies has a free parameter: The initial values for $\alpha^0$. These are used to define the habitual tendency, as well as the hard-coded prior over Flanker distractors (see Methods).

On the higher level, the context elicits a categorical observation $\mathbf{o}_t$ at the start of every episode encoded in the observation generation rules

$$p\left(\mathbf{o}_t | \mathbf{c}_t\right) = \prod_{o,c} w_{oc}^{\delta_{o,\mathbf{o}_t} \delta_{c,\mathbf{c}_t}} \tag{9}$$

where off diagonal elements of $w_{oc}$ encode context observation uncertainty, which was used to simulate effects of the CTI in the main text (Section Cue–target interval). The prior over contexts

$$p\left(\mathbf{c}_t\right) = \sum_{\mathbf{c}_{t-1}} p\left(\mathbf{c}_t | \mathbf{c}_{t-1}\right) p\left(\mathbf{c}_{t-1} | \mathbf{o}_{t-1}, \mathbf{r}_{t-1}, \pi_{t-1}\right) \tag{10}$$

is defined as the posterior predictive distribution based on the previous episode $t-1$, where a posterior over contexts can be calculated based on the context observation $\mathbf{o}_{t-1}$, rewards $\mathbf{r}_{t-1}$ and behavior $\pi_{t-1}$ of the last episode $t-1$, and a context transition probability

$$p\left(\mathbf{c}_t|\mathbf{c}_{t-1}\right) = \prod_{c',c} q_{c'c}^{\delta_{c',\mathbf{c}_t}\delta_{c,\mathbf{c}_{t-1}}} \tag{11}$$

Note that the diagonal elements of $q_{c'c}$ can be interpreted as an agent's assumption about context stability, while the off diagonal elements encode the agent's expectation of a context change. These were used in the main text to simulate the RSI curve.

For an even more detailed introduction to this type of agent and the generative model refer the interested reader to the article where the prior-based control model was originally introduced (Schwöbel et al., 2021).

#### Inference

At each time step $k$ during an episode $t$, an agent uses this generative model to infer hidden variables in the generative model, such as future states $\mathbf{s}_{k+1:K}$ and rewards $\mathbf{r}_{k+1:K}$, the context $\mathbf{c}_t$, and which policy $\pi$ to take, given observed past rewards $\mathbf{r}_{1:k}$, states $\mathbf{s}_{1:k}$, and context cue $\mathbf{o}_t$. This corresponds to calculating the posteriors

$$q\left(\mathbf{s}_{k'}\right) \approx p\left(\mathbf{s}_{k'}|\mathbf{r}_{1:k}, \mathbf{s}_{1:k}, \mathbf{o}_t\right) \tag{12}$$

$$q\left(\mathbf{r}_{k'}\right) \approx p\left(\mathbf{r}_{k'}|\mathbf{r}_{1:k}, \mathbf{s}_{1:k}, \mathbf{o}_t\right) \tag{13}$$

$$q\left(\mathbf{c}_t\right) \approx p\left(\mathbf{c}_t|\mathbf{r}_{1:k}, \mathbf{s}_{1:k}, \mathbf{o}_t\right) \tag{14}$$

$$q\left(\pi\right) \approx p\left(\pi|\mathbf{r}_{1:k}, \mathbf{s}_{1:k}, \mathbf{o}_t\right) \tag{15}$$

over the hidden variables given the observed variables. Additionally, the agent infers model parameters based on past observations using the posteriors

$$q\left(\theta\right) \approx p\left(\theta|\mathbf{r}_{1:k}, \mathbf{s}_{1:k}, \mathbf{o}_t\right) \tag{16}$$

$$q\left(\phi\right) \approx p\left(\phi|\mathbf{r}_{1:k}, \mathbf{s}_{1:k}, \mathbf{o}_t\right) \tag{17}$$

which enables the agent to learn the parameters $\theta$ of the prior over policies and the parameters $\phi$ of the reward rules.

In practice, it is often not feasible to use exact Bayesian inference, so we assume the agent uses approximate variational inference with the approximate distribution

$$q\left(\mathbf{r}_{k+1:K}, \mathbf{s}_{k+1:K}, \pi, \mathbf{c}_t, \theta, \phi\right) = q\left(\pi\right) q\left(\mathbf{c}_t\right) q\left(\theta\right) q\left(\phi\right) q\left(\mathbf{r}_{k+1:K}, \mathbf{s}_{k+1:K}|\pi\right) \tag{18}$$

where we assume a mean-field approximation except for the states and rewards of the MDP, where $q\left(\mathbf{r}_{k+1:K}, \mathbf{s}_{k+1:K}|\pi\right)$ follows the Bethe approximation, see (Schwöbel, Kiebel, & Marković, 2018; Schwöbel et al., 2021).

The approximate beliefs $q$ are found at the minimum of the variational free energy

$$\begin{aligned} q* &= \arg\min_q F\left[q\right] \\ &= \arg\min_q D_{KL}\left[q\left(\mathbf{r}_{k+1:K}, \mathbf{s}_{k+1:K}, \pi, \mathbf{c}_t, \theta, \phi\right)|p\left(\mathbf{r}_{1:K}, \mathbf{s}_{1:K}, \phi, \pi, \theta, \mathbf{c}_t, \mathbf{o}_t\right)\right] \end{aligned} \tag{19}$$

which is the Kullback-Leibler divergence between the approximate posterior $q$ and the generative model $p$.

#### Belief updates

In this section, we show the belief updating equations that minimize the variational free energy. For a full derivation of these equations we refer the interested reader to our previous work (Schwöbel et al., 2018, 2021).

The approximate posterior over MDP variables – future states $\mathbf{s}_{k'}$ and future rewards $\mathbf{r}_{k'}$ – is given by the belief propagation message passing rules (Schwöbel et al., 2018) as

$$\begin{aligned}
q\left(\mathbf{r}_{k'},\mathbf{s}_{k'}|\pi,\mathbf{c}_t\right) &= \frac{p\left(R|\mathbf{r}_{k'}\right)p'\left(\mathbf{r}_{k'}|\mathbf{s}_{k'},\mathbf{c}_t\right)m_\pi^{k'+1}\left(\mathbf{s}_{k'}|\mathbf{c}_t\right)m_\pi^{k'-1}\left(\mathbf{s}_{k'}|\mathbf{c}_t\right)}{Z_{k'}^\pi} \\
q\left(\mathbf{r}_{k'}|\pi,\mathbf{c}_t\right) &= \frac{p\left(R|\mathbf{r}_{k'}\right)m_\pi^{k'}\left(\mathbf{r}_{k'}|\mathbf{c}_t\right)}{Z_{k'}^\pi} \\
q\left(\mathbf{s}_{k'},\mathbf{s}_{k'-1}|\pi,\mathbf{c}_t\right) &= \frac{p\left(\mathbf{s}_{k'}|\mathbf{s}_{k'-1},\pi\right)}{Z_{k',k'-1}^\pi}m_r^{k'-1}\left(\mathbf{s}_{k'-1}\right)m_r^{k'}\left(\mathbf{s}_{k'}\right)m_\pi^{k'+1}\left(\mathbf{s}_{k'}|\mathbf{c}_t\right)m_\pi^{k-2}\left(\mathbf{s}_{k'-1}|\mathbf{c}_t\right) \\
q\left(\mathbf{s}_{k'}|\pi,\mathbf{c}_t\right) &= \frac{m_\pi^{k'+1}\left(\mathbf{s}_{k'}|\mathbf{c}_t\right)m_\pi^{k'-1}\left(\mathbf{s}_{k'}|\mathbf{c}_t\right)}{Z_{k'}^\pi}
\end{aligned} \tag{20}$$

using the messages

$$\begin{aligned}
m_r^{k'}\left(\mathbf{s}_{k'}|\mathbf{c}_t\right) &= \sum_{\mathbf{r}_{k'}}p\left(R|\mathbf{r}_{k'}\right)p'\left(\mathbf{r}_{k'}|\mathbf{s}_{k'},\mathbf{c}_t\right), \\
m_\pi^{k'}\left(\mathbf{r}_{k'}|\mathbf{c}_t\right) &= \sum_{\mathbf{s}_{k'}}p'\left(\mathbf{r}_{k'}|\mathbf{s}_{k'},\mathbf{c}_t\right)m_\pi^{k'+1}\left(\mathbf{s}_{k'}|\mathbf{c}_t\right)m_\pi^{k'-1}\left(\mathbf{s}_{k'}|\mathbf{c}_t\right), \\
m_\pi^{k'+1}\left(\mathbf{s}_{k'}|\mathbf{c}_t\right) &= \frac{1}{Z'_{k',\pi}}\sum_{\mathbf{s}_{k'+1}}p\left(\mathbf{s}_{k'+1}|\mathbf{s}_{k'},\pi\right)m_r^{k'+1}\left(\mathbf{s}_{k'+1}|\mathbf{c}_t\right)m_\pi^{k'+2}\left(\mathbf{s}_{k'+1}|\mathbf{c}_t\right), \\
m_\pi^{k'-1}\left(\mathbf{s}_{k'}|\mathbf{c}_t\right) &= \frac{1}{Z''_{k',\pi}}\sum_{\mathbf{s}_{k'-1}}p\left(\mathbf{s}_{k'}|\mathbf{s}_{k'-1},\pi\right)m_r^{k'-1}\left(\mathbf{s}_{k'-1}|\mathbf{c}_t\right)m_\pi^{k'-2}\left(\mathbf{s}_{k'-1}|\mathbf{c}_t\right),
\end{aligned} \tag{21}$$

where

$$\ln p'\left(\mathbf{r}_{k'}|\mathbf{s}_{k'},\mathbf{c}_t\right) = \int d\phi q\left(\phi\right)\ln p\left(\mathbf{r}_{k'}|\mathbf{s}_{k'},\phi,\mathbf{c}_t\right). \tag{22}$$

The posterior over policies for each context is

$$q\left(\pi|\mathbf{c}_t\right) \propto p'\left(\pi|\mathbf{c}_t\right)\exp\left(-F\left(\pi|\mathbf{c}_t\right)\right) \tag{23}$$

where $p'\left(\pi|\mathbf{c}_t\right)$ is the marginalized prior over policies, and $F\left(\pi|\mathbf{c}_t\right)$ is the policy-specific free energy in a given context (see (Schwöbel et al., 2018)). The policy-specific free energy encodes how surprising and rewarding policies will be in this context.

The posterior over the parameters $\theta$ of the prior over policies is only updated at the end of an episode, when a policy has been executed fully. The update is given as

$$\begin{aligned}
q\left(\theta\right) &= \frac{1}{B\left(\alpha^t\right)}\prod_{l,n}\theta_{ln}^{\alpha_{ln}^t-1} \\
\alpha_{ln}^t &= \lambda\alpha_{ln}^{t-1} + q\left(\pi=l|\mathbf{c}_t=n\right)q\left(\mathbf{c}_t=n\right)+1-\lambda
\end{aligned} \tag{24}$$

and is itself again a Dirichlet distribution with updated pseudo counts $\alpha^t$. They are updated based on which policy was chosen in which context, and are discounted by the forgetting factor $\lambda$. $\lambda$ was set to 1 in the grid world as well as in the task switching task, and to 0.9 in the Flanker task.

The posterior over the parameters $\phi$ of the outcome rules is given as

$$\begin{aligned} q(\phi) &= \frac{1}{B(\beta^t)} \prod_{i,j,n} \phi_{ijn}^{\beta_{ijn}^t - 1} \\ \beta_{ijn}^t &= \beta_{ijn}^{t-1} + q(\mathbf{c}_t = n) \sum_{m=1}^{k} \delta_{i,\mathbf{r}_m} \delta_{j,\mathbf{s}_m} \end{aligned} \tag{25}$$

where the concentration parameters $\beta^t$ are updated by the state-reward pairs that were perceived so far in the current episode.

Lastly, the posterior over contexts is

$$\begin{aligned} \ln q(\mathbf{c}_t) &\propto \ln p'(\mathbf{c}_t) + \ln p\left(\mathbf{o}_t|\mathbf{c}_t\right) - D_{KL}\left[q\left(\pi|\mathbf{c}_t\right)|p'\left(\pi|\mathbf{c}_t\right)\right] - \sum_{\pi} q\left(\pi|\mathbf{c}_t\right) F(\pi, \mathbf{c}_t) \\ q(\mathbf{c}_t) &\propto p'(\mathbf{c}_t) p\left(\mathbf{o}_t|\mathbf{c}_t\right) \exp\left(-F(\mathbf{c}_t)\right) \end{aligned} \tag{26}$$

with context-specific free energy $F(\mathbf{c}_t)$. Note, that we set

$$p'(\mathbf{c}_t) = \sum_{\mathbf{c}_{t-1}} q(\mathbf{c}_{t-1}) p\left(\mathbf{c}_t|\mathbf{c}_{t-1}\right) \tag{27}$$

as a prior over the current context based on the context transition probabilities $p\left(\mathbf{c}_t|\mathbf{c}_{t-1}\right)$.

#### Task parameters

We shown in Table 1 the parameters that were set for the simulations of the three tasks in the main text. For notational clarity we use $q_{cc} = p\left(\mathbf{c}_t = c|\mathbf{c}_{t-1} = c\right)$ for the probability to stay in a context, see also Eq. 11; and $w_{oc} = p\left(\mathbf{o}_t|\mathbf{c}_t\right)$ for the off-diagonal elements of the context observation generation probability which encodes the context observation uncertainty, see Eq. 9.

| | $q_{cc}$ | $w_{oc}$ | $\alpha^0$ | $\lambda_\pi$ |
|---|---|---|---|---|
| Grid world $h = 0.001$ | 97% | – | 1000 | 0 |
| Grid world $h = 1$ | 97% | – | 1 | 0 |
| Flanker | 95% | 0% | 100 1<br>1 100 | 0.1 |
| Task switching | 92% | 0.9% | 1000 1000<br>1000 1000 | 0 |

**Table 1.** Parameter settings for the simulated agents in the three experiments below. The columns show the parameter types: $q_{cc}$ is the context stay probability (see Eq. 11), $w_{oc}$ is the cue uncertainty (see Eq. 9), $\alpha^0$ are the initial pseudo counts in the Dirichlet prior (see Eq. 8) and $\lambda$ is the forgetting factor (see Eq. 24). The rows show the task types. In the grid world we show two agents, a weak prior learner ($h = 0.001$) and a strong prior learner ($h = 1$). The context stability probability was set to 97% in both cases, and there were no context cues so there is no context cue uncertainty. The prior over policies in this case has shape $n_\pi \times n_c = 81 \times 20$, and for each entry the initial pseudo counts were set the same. Hence the table shows only one value. In the flanker task, there are two actions and two contexts, hence the initial pseudo counts form a $2 \times 2$ matrix. They were preset to favor action 1 in context 1 and action 2 in context 2. Note that in this task, the pseudo counts are also subject to a small forgetting factor $\lambda = 0.1$ in each trial, which was set to zero, i.e. no forgetting, in all other tasks. In the task switching task, there are also $2 \times 2$ initial pseudo counts, but they were set to the same high value so that the prior does have little to no influence on action selection.
