## Supplement 2, flanker details for "Joint modeling of choices and reaction times based on Bayesian contextual behavioral control"

### Supplementary file 2: Flanker features

To simulate main effects of the Flanker Task in the main text, the interplay between context and prior over actions is of key importance. Namely agents were set up so that the flankers elicit flanker–response associations, which were encoded as context–prior associations that favor the response associated with the flanker in the quickly executable prior over actions. In order to underline how important this interplay is for both flanker effects, the conditional accuracy function as well as the Gratton effect, we will show here three different modified agents with different parts of this machinery switched off.

Firstly, we simulated 50 agents with only the learning of the prior switched off (Figure 1). Here, the agents start with pre-existing flanker–response associations but do not adapt their strength according to a learning rule, as was done in the main text. In this case, the conditional accuracy function (Figure 1A) shows that in congruent trials, agents are always correct, and due to being very certain of being correct, are also always very fast to respond. However in incongruent trials, agents are almost always wrong and choose the flanker response instead of the one indicated by the stimulus in the middle. This is visibly different from the typical conditional accuracy function shown in the main text, which is usually measured in flanker tasks (Gratton, Coles, & Donchin, 1992; White, Brown, & Ratcliff, 2012). Due to the learning of the prior missing in the agents, the sequential effects usually found in the Gratton effect are also not present, see Figure 1B. Instead, responses in congruent trials are simply faster than in incongruent trials, but the previous trial type does not affect the response time.

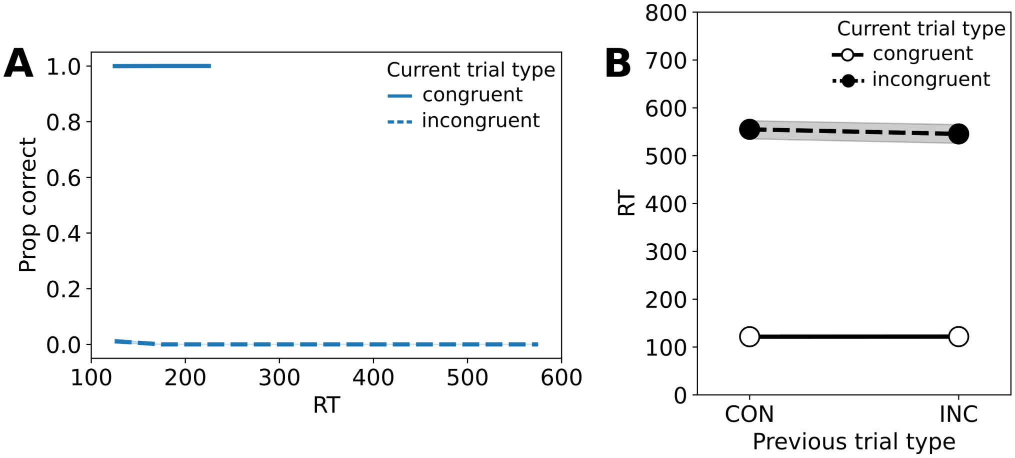


**Fig 1. Flanker effects under switched off prior learning. A** Conditional accuracy function, **B** sequential effects. Solid and dashed lines indicate congruent and incongruent trials, respectively.

Next, we show that these pronounced flanker–response associations in the prior are important to the flanker results. Hence we exchanged the pre-set pronounced prior with a flat one in another 50 agents (Figure 2). Here, flankers are ineffective, as they do not elicit an automatic urge to choose the response indicated by the flanker and encoded in the prior. As a consequence, the accuracy of choices in congruent and incongruent trials does not differ, and agents are mostly correct in both trial types, see Figure 2A. Similarly, the reaction times do not differ for congruent and incongruent responses as shown in Figure 2B. In fact, reaction times for both in this scenario are slowed down, as the speeding up effect of the prior is removed.

Lastly, we illustrate that simulating a context explicitly is important for the flanker task. Hence we set up 50 agents using only one known context (Figure 3). In this scenario, it is not possible to have pre-determined flanker–response associations, as

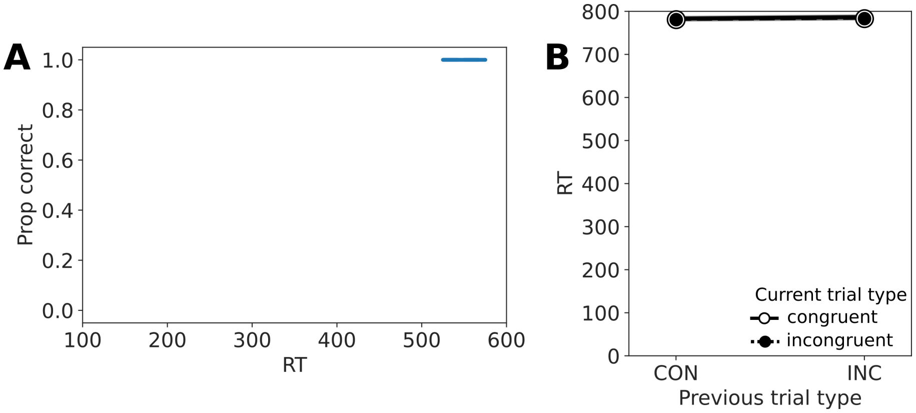


**Fig 2. Flanker effects with a flat prior. A** Conditional accuracy function, **B** sequential effects. Solid and dashed lines indicate congruent and incongruent trials, respectively.

there would need to be two contexts for there to be association with two responses. Similarly to the case with the flat prior, agents do not process the flankers any more, and hence are mostly correct in congruent as well as incongruent trials, see Figure 3A. Additionally, reaction times in both cases are the same (Figure 3B). However, the reaction times in this scenario are a bit faster than in the flat prior scenario shown in Figure 2B, because one context means "one task", similar to the single task simulations in the Task switching section in the main text, where reaction times are also reduced compared to a two context scenario.

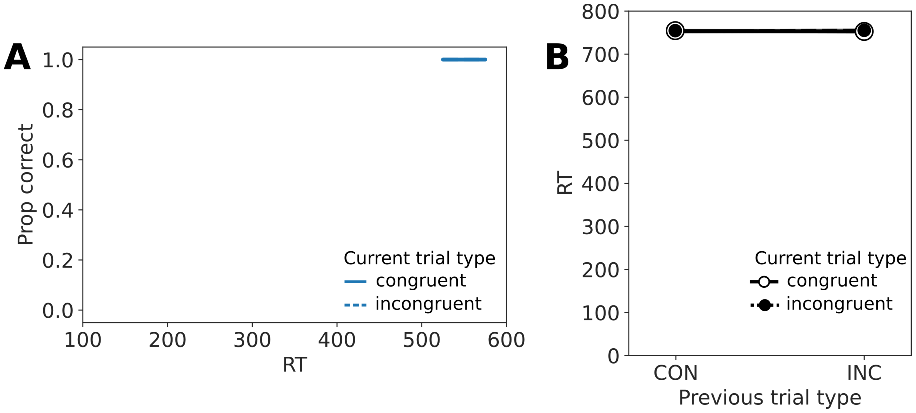


**Fig 3. Flanker effects with only one context. A** Conditional accuracy function, **B** sequential effects. Solid and dashed lines indicate congruent and incongruent trials, respectively.
