## Supplement 3, task switching details for "Joint modeling of choices and reaction times based on Bayesian contextual behavioral control"

### Supplementary file 3: Task switching features

In the simulated task switching task, the prior was set to flat and learning was negligible, so changing features of the prior learning will not affect the resulting agent behavior. Due to the contextual nature of the task however, the context-specific learning of action–outcome contingencies, i.e. the task set, is a key feature only due to which an agent is able to learn two different tasks and switch between them. In order to show this, we have simulated 50 agents who only learn within one context, and left the rest of the task setup as it is in the main text. The resulting reaction times and accuracies are shown in Figure 1.

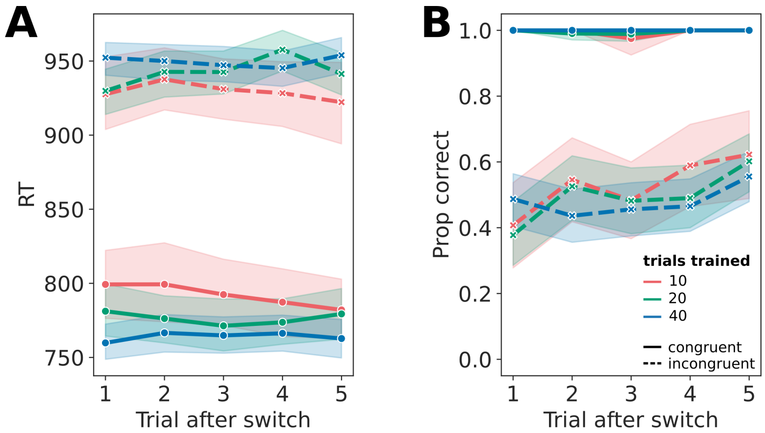


**Fig 1. Reaction time and accuracy as a function of trial after switch. A** Reaction times of 50 simulated agents who use only one context in a task switching task. Solid lines indicate congruent trials, and dashed lines indicate incongruent trials. The colors indicate how many trials have been trained before. **B** Proportion of correct choices, i.e. accuracies in the same experiment from **A**, color scheme also as in **A**.

When task set 1 is active during the experiment, e.g. a letter task where it needs to be indicated whether a letter is a vowel or a consonant, the agent learns which letter stimulus corresponds to which response key, e.g. left or right. When task set 2 is active, e.g. a number task where it is to be indicated whether a number is even or odd, the agent learns which key indicates the correct response. In the agent setup in the main text, agents learn these two task sets as different action–outcome contingencies in two distinct contexts. They can infer which of the two is active by interpreting the context cue and choosing a response according to the currently active task set.

If agents however only have one context available in which they learn both task sets and action–outcome contingencies, no task set specific representation can be learned. As a consequence, agents learn to associate congruent stimuli with one specific action, e.g. vowel and even with the right key, and consonant and odd with the left key. These contingencies can be learned well, as can be seen in the high choice accuracy for congruent trials in Figure 1B. Accordingly, and since agents are fairly certain of their assessment which key is correct, the corresponding reaction times are low for congruent trials, see Figure 1A.

However, in incongruent trials, agents are seemingly randomly rewarded for both responses, as the agent cannot understand that there are two different task sets. Hence the random choices in incongruent trials in Figure 1B, and due to being very uncertain about the correct response, very long reaction times in Figure 1A. If there is an incongruent trial in the first trial after a switch, agents start out with choosing the correct action with a bit below chance probability, as they learned the opposite contingencies in the previous run. This probability slowly increases with each consecutive trial in the same task set, as the agents slowly unlearn the previous contingencies and learn the ones of the current task set.

That a learning of contingencies and rewards is still happening is evidenced by the fact that even in this setting, agents become quicker in the congruent trials the more trials they have trained in the experiment in the past (Figure 1A). However, the opposite happens in the incongruent trials where agent become slower and slower during the course of the experiment.

The RSI and CSI effects in the main text were simluated using variations of parameters that agents use for context inference, namely the expected context change probability and the context cue uncertainty. If there is only one context, the context change probability is always zero, as there is no second context that the agent could change into. Similarly, since there is only one context, both task cues will be associated with the same context, and there cannot be any context cue uncertainty. Hence, in the agent setting with only one context, it is not possible to simulate either inter-trial effects like the RSI, or cue presentation effects like the CSI which is why we cannot show adapted versions of these results here.
